# Spatial multi-omics defines a shared glioblastoma infiltrative signature at the resection margin

**DOI:** 10.1101/2024.11.05.621879

**Authors:** Balagopal Pai, Susana Isabel Ramos, Wan Sze Cheng, Tanvi Joshi, Gabrielle Price, Jessica Tome-Garcia, German Nudelman, Sanjana Shroff, Kristin Beaumont, Yong Raymund, Robert Sebra, Elena Zaslavsky, Nadejda Mincheva Tsankova

## Abstract

Glioblastoma (GBM) remains an untreatable disease. Understanding GBM’s infiltrative biology at the resection margin is limited, despite causing disease recurrence and progression. To address this, we generated a high-throughput single-nucleus (sn)RNA-seq and snATAC-seq multi-omic dataset from six tumors with distinct genomic drivers and combined it with spatial transcriptomics to characterize the unique molecular phenotype of GBM near the margin. By contrasting GBM-specific biology in matching “Core” vs. “Margin” dissections, we define unique, shared “GBM infiltration” and chromatin accessibility signatures near the margin. We prioritize *EGFR* as a top differentially expressed and accessible “Margin” marker across GBM subtypes, show its dynamic expression along a core-to-margin infiltration trajectory, and validate its role in migration through CRISPR/Cas9 deletion in two patient-derived models. ChIP-seq studies furthermore corroborate preferential TEAD1 binding at EGFR’s accessible regulatory elements. This validated multi-omic dataset enables further studies into tumor and microenvironment biology in the context of residual GBM disease.

## Introduction

Glioblastoma (GBM), the most common malignant brain tumor in adults, continues to carry a dismal prognosis despite aggressive therapy. An important and poorly understood contributor to treatment resistance and progression in GBM is the diffusely infiltrative spread of tumor cells, a shared malignant behavior regardless of tumor heterogeneity [1-3]. Standard-of-care treatment in patients with GBM involves gross-total resection of the tumor “Core,” the contrast-enhancing mass on MRI, which histologically displays high tumor density and necrosis; followed by chemoradiation targeting residual tumor at the “Margin,” a poorly defined MRI area that consists of scattered GBM cells infiltrating into the patient’s brain [4]. Peripherally infiltrating GBM cells away from the tumor core adapt to unique microenvironmental pressures [3, 5-7]. Such adaptations have been attributed to plasticity within glioma stem cell-like (GSC) populations, which are highly infiltrative and linked to tumor progression [3, 8-10].

Our understanding of GBM biology at the infiltrative margin remains limited, in part due to preferential use of patient-derived models derived from “Core” specimens, limited supratotal resection surgeries [11] with available “Margin” tissue, and the technical difficulty of isolating rare GBM cells admixed with mostly normal brain. Yet, it is the residual disease at the margin that is treated with chemoradiation and eventually causes recurrence [3]. Current single-cell technologies enable isolation and profiling of rare tumor populations with high resolution and spatiotemporal context. While such high-throughput efforts initially focused on GBM heterogeneity in highly-cellular “Core” tissue [12-21], notable studies have begun to characterize tumor [22-33] and tumor microenvironment (TME) [34, 35] biology at the infiltrative periphery.

This study builds on the latter efforts by creating a high-throughput multi-omic dataset representative of GBM biology at the most distal region of tumor infiltration near the left-behind “Margin,” towards understanding better the unique biology of residual disease. To this end, we isolated and profiled “Core” and “Margin” GBM cells from six surgical tumors with diverse mutational drivers. Using spatial transcriptomics (ST) along with matched single-nucleus (sn)RNA-seq and snATAC-seq, we define a unique “GBM infiltration” signature at the Margin, shared across tumors with diverse genomic drivers. We experimentally validate one of the top GBM-Margin genes, *EGFR*, as a driver of GBM migration in patient-derived models, and show preferential occupancy of the targetable GSC regulator, TEAD1 [36-38], at its locus. This high-quality, genomically diverse multi-omic dataset serves as an important new resource for the community to scrutinize further residual disease biology at the infiltrative margin.

### Spatial multi-omic dataset resolves distinct cell types at the GBM margin

To better characterize the unique biology of infiltrative tumor cells near the resection margin, we performed multi-omics on GBM tissue dissections from contrast-enhancing tumor “Core” and from distal FLAIR-hyperintense regions near the surgical “Margin” of rare supratotal resections (Fig. 1a-b). Neuropathological evaluation confirmed a hypercellular tumor within the “Core” with characteristic palisading necrosis and microvascular proliferation and predominantly non-neoplastic brain parenchyma within the “Margin”, with scattered glioma cells infiltrating as single cells or forming satellitosis (Fig. 1c). snRNA-seq and snATAC-seq were performed from the same “Core” and “Margin” nuclei dissociates in six patient-derived glioblastoma (CNS WHO grade 4) specimens, maximizing genomic diversity (Fig. 1d, Extended Data 1). Of note, GBM2 was unusual in its co-presence of EGFR amplification and IDH mutation, with this and other such tumors displaying aggressive, GBM-like behavior with short survival [39, 40], justifying inclusion in our cohort.

**Figure 1:**
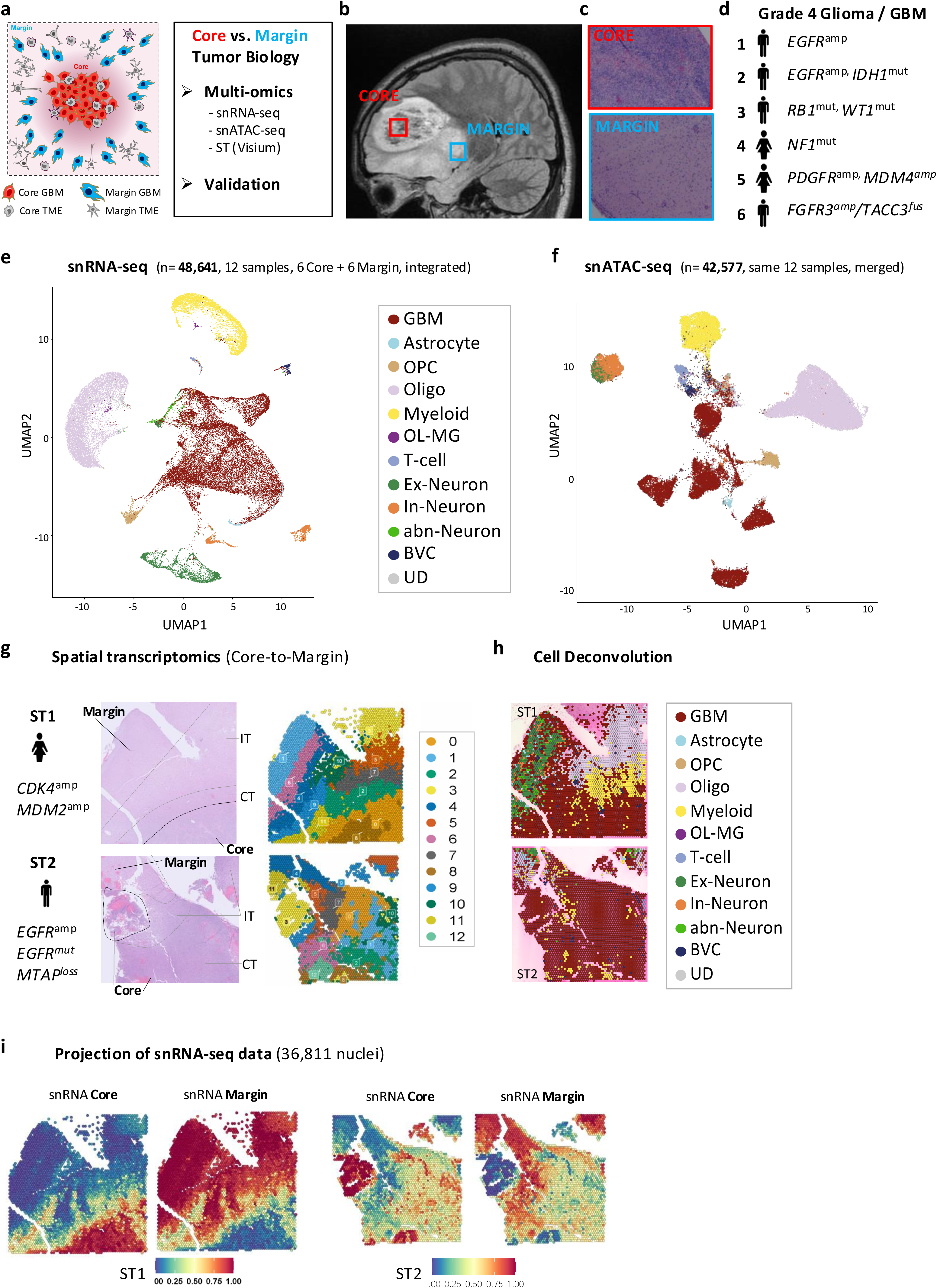
Spatial multi-omic profile of GBM biology at Core and Margin. **a.** Schematic of GBM and tumor microenvironment (TME) composition at Core (red) and Margin (blue) and study workflow. **b.** Preoperative T1-weighted Magnetic Resonance Image (MRI) from patient GBM2 showing representative tumor Core and Margin regions targeted for the study. **c.** Hematoxylin and eosin (H&E)-stained images from GBM2, showing representative histology at Core and Margin. **d.** Cohort of diverse tumor specimens profiled in study showing main genomic drivers and patient sex. **e.** UMAP of snRNA-seq data from all GBM samples, integrated, colored by cell type. See also Supplementary Figs. 1, 3, 4. **f.** UMAP of snATAC-seq data from all GBM samples, merged, colored by cell type. See also Supplementary Fig. 2. **g.** Spatial transcriptomics data from two additional GBM samples, ST1 and ST2, capturing Core-to-Margin transition. Shown are main genomic drivers, sex, and neuropathology-based regional annotations (left) and corresponding SpatialDimPlot clusters (right). IT = infiltration tumor, CT = cellular tumor. See also Supplementary Fig. 5a. **h.** SpatialDimplot of cell type-enriched ROI inferred from Seurat-based cell type deconvolution using snRNA-seq data. See also Supplementary Fig. 5b-d. **i.** SpatialFeaturePlots of ST1 (left) and ST2 (right) showing enrichment of projected snRNA-seq “Core” and “Margin” data to expected spatial regions (confident dissections only, GBM1+GBM2+GBM4+GBM6). See also Supplementary Fig. 6.

All 12 (six “Core” and six “Margin”) sequenced samples were mapped to hg38 and underwent standard quality assessment (see Methods) (Extended Data 2). After filtering, snRNA-seq data (48,641 nuclei) were SCT-normalized and integrated using Seurat [41, 42], and then used for initial annotation of cell types (Fig. 1e). Despite genomic variability, the data integrated well, with each GBM sample contributing to each cluster (Supplementary Fig. 1a-b). Clusters were annotated using canonical markers for the following cell-types: GBM, astrocytes, oligodendrocyte progenitor cells (OPCs), myelinating oligodendrocytes, Myeloid, T-cell, excitatory and inhibitory neurons, and blood vessels cells (BVC) (Fig 1e, Supplementary Fig. 1c). We also identified a previously described *MBP*+*CX3CR1*+ inflammatory oligodendroglial population (OL-MG) [43-45] and a poorly defined neuronal population with aberrant phenotype (abn. neuron) (Fig. 1e, Supplementary Fig. 1c). Cell type annotations were confirmed via projections from an external neurotypic adult cortex snRNA-seq dataset [46] (Supplementary Fig. 1d).

To correlate gene expression with open chromatin and transcription factor (TF) motif activity in each GBM region, we generated matching snATAC-seq data from the same 12 GBM samples (n=42,577 nuclei after filtering) (Fig. 1f, Extended Data2). We visualized snATAC-seq data using both a merged object (Fig. 1f, Supplementary Fig. 2a), and integrated using Harmony (Supplementary Fig. 2b) [47, 48]. Both showed integration of non-neoplastic populations with expected batch effects seen in the tumor component when merged. Subsequent snATAC-seq visualization was done on merged objects, to better discern tumor-specific effects. Cross-platform integration (see Methods) demonstrated robust correspondence of snRNA-seq and snATAC-seq datasets (Supplementary Fig. 2c) and enabled confident annotation of ATAC cell type clusters (Supplementary Fig. 2d).

To confirm malignant cell designation, we inferred copy number alterations in our snRNA-seq dataset using InferCNV [12], using both external cortex [46] (Supplementary Fig. 3) and internal non-neoplastic populations (Supplementary Fig. 4a) as references. Inference scores for chromosome 7 gain (+7) and chromosome 10 loss (-10), the most frequent and defining cytogenetic alterations in GBM [2, 40], were high in tumor clusters and negligible elsewhere (Supplementary Fig. 4b). A combined score including +7/-10 CNV and GSC markers increased tumor specificity further (Supplementary Fig. 4c-d). Cell type proportions analysis of snRNA-seq data showed expected predominance of tumor cells in “Core”, along with some myeloid and BVCs; and predominance of non-neoplastic cell types in the “Margin” (Supplementary Fig. 1e). Of note, GBM3 showed a small cluster of tumor cells in Core, and GBM5 showed similar tumor cell proportions in “Core” and “Margin” (Supplementary Fig. 1e, Supplementary Fig. 4a), suggesting that tissue dissection in these two samples may have been suboptimal.

To provide additional spatial context onto dissociated snRNA-seq Core/Margin, we also generated high-quality ST data for two additional GBM samples derived from supratotal gross total resection (4010 ST1 and 3111 ST2 spots after filtering) (Fig. 1g, Supplementary Fig. 5a, Extended Data2). Distinct anatomical regions, including necrotic Core, cellular tumor (CT), infiltrative tumor (IT), and distal leading edge (Margin) were annotated by an experienced neuropathologist before and after ST processing (Fig. 1g, left). ST1 demonstrated clear transition in tumor density from “Core” to “Margin” on histology, corresponding SpatialDimPlot (Fig. 1g, right) and UMAP cluster-based separation (Supplementary Fig. 5a). ST2 had a less organized structure, but histological annotations still correlated to unbiased clustering of spatial data (Fig. 1g). To independently identify ST regions-of-interest (ROI) enriched in tumor cells, we: a) calculated +7/-10 inferred CNV expression (Supplementary Fig. 5b); b) projected snRNA-seq GBM cell-type signatures onto ST data (Supplementary Fig. 5c); and c) performed cell deconvolution using Seurat [42] (Fig. 1h) and RCTD [49] (Supplementary Fig. 5d). This enabled estimation of cell-type specific contribution to each spatial region and extraction of cell-type enriched ROI. Deconvolution showed expected laminar-pattern of neuron-enriched ROI at the infiltrated neocortex near the “Margin”, oligodendroglia-enriched ROI within infiltrated white matter (IT), and high tumor density within “Core” and “CT” (Fig. 1h, Supplementary Fig. 5c-d). Importantly, scattered GBM-enriched ROI were also recovered within “IT” and “Margin” regions.

Finally, to confirm “Core” and “Margin” designation from our dissections, we projected dissociated snRNA-seq data onto ST1 and ST2 (Fig. 1i), including separately for each GBM sample (Supplementary Fig. 6). GBM1, GBM2, GBM4 and GBM6 snRNA-seq projections matched to ST “Core” and “Margin”, providing high confidence in our dissections. In contrast, GBM3 did not match and GBM5 showed only partial match (Supplementary Fig. 6), justifying their removal from downstream “Margin”-based analyses (Fig. 2a-b, Extended Data 3).

**Figure 2:**
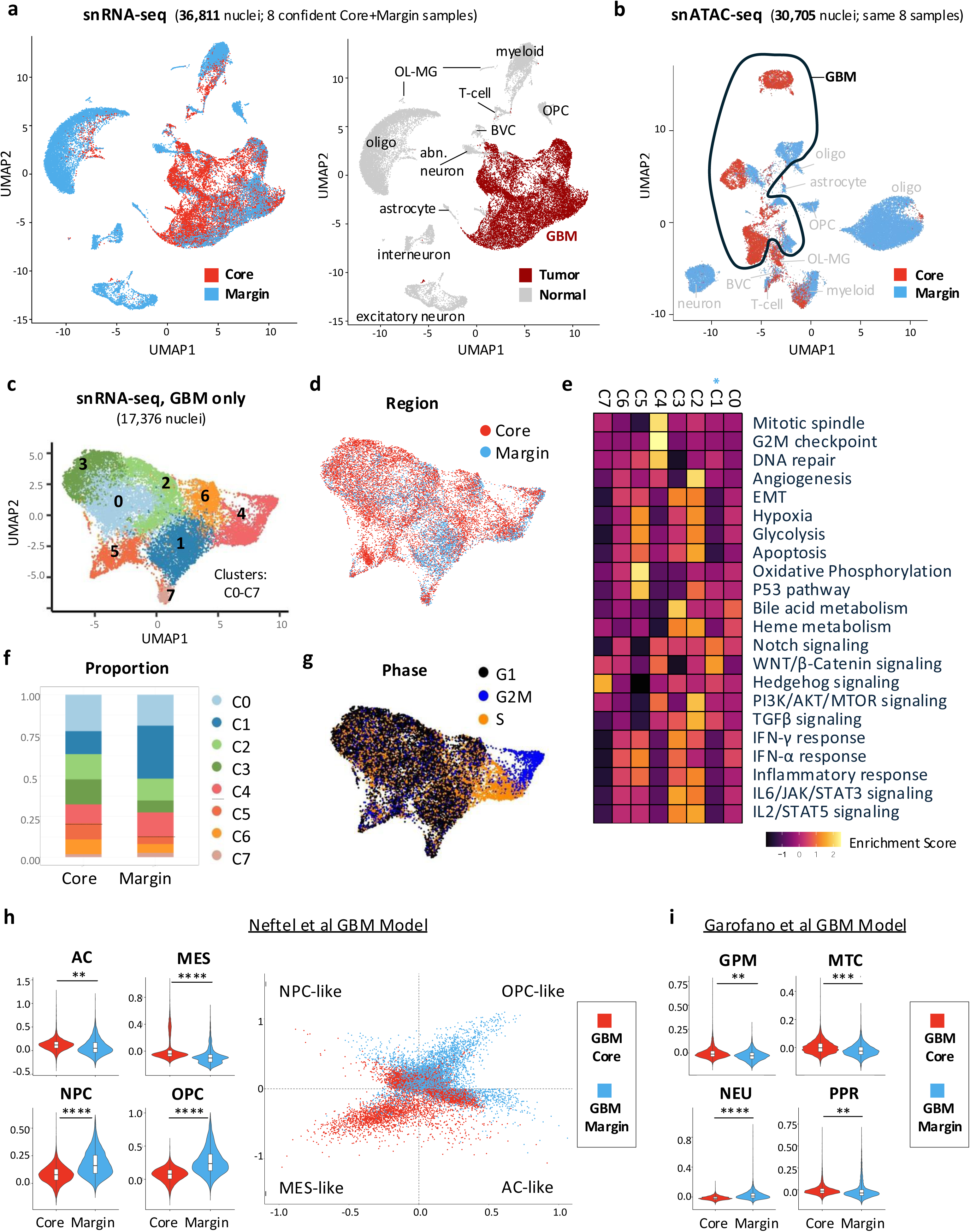
Analysis of snRNA-seq GBM Core and Margin data using pre-defined models. **a.** UMAP of snRNA-seq GBM data from confident Core and Margin dissections, colored by region (left) and by tumor status (right). See also Supplementary Fig. 1f. **b.** UMAP of snATAC-seq GBM data from confident Core and Margin dissections, colored by region. See also Supplementary Fig. 2a. **c.** UMAP of GBM-subset snRNA-seq data from confident Core and Margin dissections, colored by cluster number. **d.** UMAP of GBM-subset snRNA-seq data from confident Core and Margin dissections, colored by region. **e.** Heatmap of enrichment scores for select Hallmark terms in GBM-subset clusters. **f.** Stacked bar plot of nuclei proportions (% of total) in Core and Margin for each GBM-subset cluster. **g.** UMAP of G1, G2M and S cell cycle phase in GBM-subset analysis. **h.** Violin plots (left) and two-dimensional representation (right) of Neftel et al cellular state GBM model showing meta module score enrichment in GBM-Core and GBM-Margin populations (* = p.adj < e^-100^, ** = p.adj < e^-200^, *** = p.adj < e^-300^,**** = p.adj < e^-400^, unpaired Wilcoxon rank test) **i.** Violin plots showing enrichment scores of Garofano et al pathway-based GBM model in GBM-Core and GBM-Margin populations. Glycolytic/plurimetabolic (GPM) and Mitochondrial (MTC) in the top panel, Neuronal (NEU) and Proliferative/progenitor (PPR) in the bottom panel. (* = p.adj < e^-100^, ** = p.adj < e^-200^, *** = p.adj < e^-300^, **** = p.adj < e^-400^, unpaired Wilcoxon rank test)

Overall, we generated a high-quality single-cell resolution multi-omic GBM dataset with spatial separation of tumor “Core” and “Margin”, which also captures the full repertoire of TME cell types in both tumor niches. We focused downstream analyses on GBM-specific biology at the Margin and provide the dataset as a resource to the community for further tumor and TME-focused studies.

### Unique and tumor-specific “GBM infiltration” signature at the Margin

The ability to confidently label tumor cells in our snRNA-seq/snATAC-seq dataset provided new opportunities to study pure GBM biology from “Core” and “Margin”, without non-neoplastic contaminants. A high yield of GBM-Core (n=12,621) and GBM-Margin (n=4,755) nuclei also provided sufficient computational power to perform subsequent integrated analyses by sub-clustering tumor cells only (Fig. 2c, Extended Data 3). First, we assessed tumor heterogeneity per cluster (Fig. 2c-e), noting enrichment of GBM-Core in clusters C0, C2, C3, C5 and C6; predominance of GBM-Margin in cluster C1 (Fig. 2f); and similar distribution within cycling cell cluster C4 (Fig. 2f-g). Functional enrichment analysis for Hallmark pathways captured GBM heterogeneity in our dataset, with expected enrichment for hypoxia, apoptosis, glycolysis, oxidative phosphorylation, and inflammatory response signatures in Core-enriched clusters [18, 20] (Fig. 2e, Extended Data 4). Notably, “Epithelial-to-mesenchymal transition” (EMT), previously implicated in tumor invasion [50], was also enriched in Core-high clusters, while NOTCH and WNT/Beta-catenin signaling were enriched in Margin-predominant cluster C1 (Fig. 2e). Several cell division pathways were enriched in C4, confirming that a subset of both Core and Margin GBM cells are actively proliferating [3] (Fig. 2f-g).

We next assessed how GBM “Core” and “Margin” tumor biology relates to previously established classifiers of GBM heterogeneity, including based on cellular states (Neftel et al) [16] and metabolic dependance (Garofano et al) [18]. In the Neftel et al. model, GBM-Margin cells were enriched for “OPC-like” and “NPC-like” states, whereas GBM-Core cells were more “MES-like” and “AC-like” (Fig. 2h). In the Garofano et. al. model, GBM-Margin cells were enriched for neuronal “NEU” and depleted for metabolic “MTC” and “GPM”, and for proliferative/progenitor “PPR” signatures (Fig. 2i). We also spatially profiled enrichment of known gene sets related to cell invasion and migration, taking advantage of our ST data, which captured the full transition of tumor architecture from “Core”, “CT”, “IT”, and up to distal “Margin” (Fig. 1g). We improved on prior analyses with ST-Visium by deconvolving tumor specific enrichment, to more faithfully project GBM signatures onto GBM-enriched ROI (Fig. 1h, Fig. 3a). “Hallmark-Hypoxia”, “Hallmark-EMT”, and “multicancer invasion” [51] signatures were enriched in GBM “Core” across both ST (Fig. 3b) and snRNA-seq datasets (Fig. 3c). Interestingly, recently defined “invasivity” signature in unconnected GBM populations at the tumor periphery [23, 24] as well as “cell migration” showed enrichment at the ST transition zone between Core and Margin, annotated as CT and IT (Fig. 3a-b). There was highly significant enrichment for “Cell migration” in GBM-Margin snRNA-seq populations with more modest significance for “invasivity” (Fig. 3c). Analysis of layered GBM architecture signatures derived from spatial data [28] showed significant enrichment of “Layers4-5” in GBM-Margin populations (Supplementary Fig. 7a), with non-tumor contribution within the “Layers4-5” signature potentially confounding this analysis.

**Figure 3:**
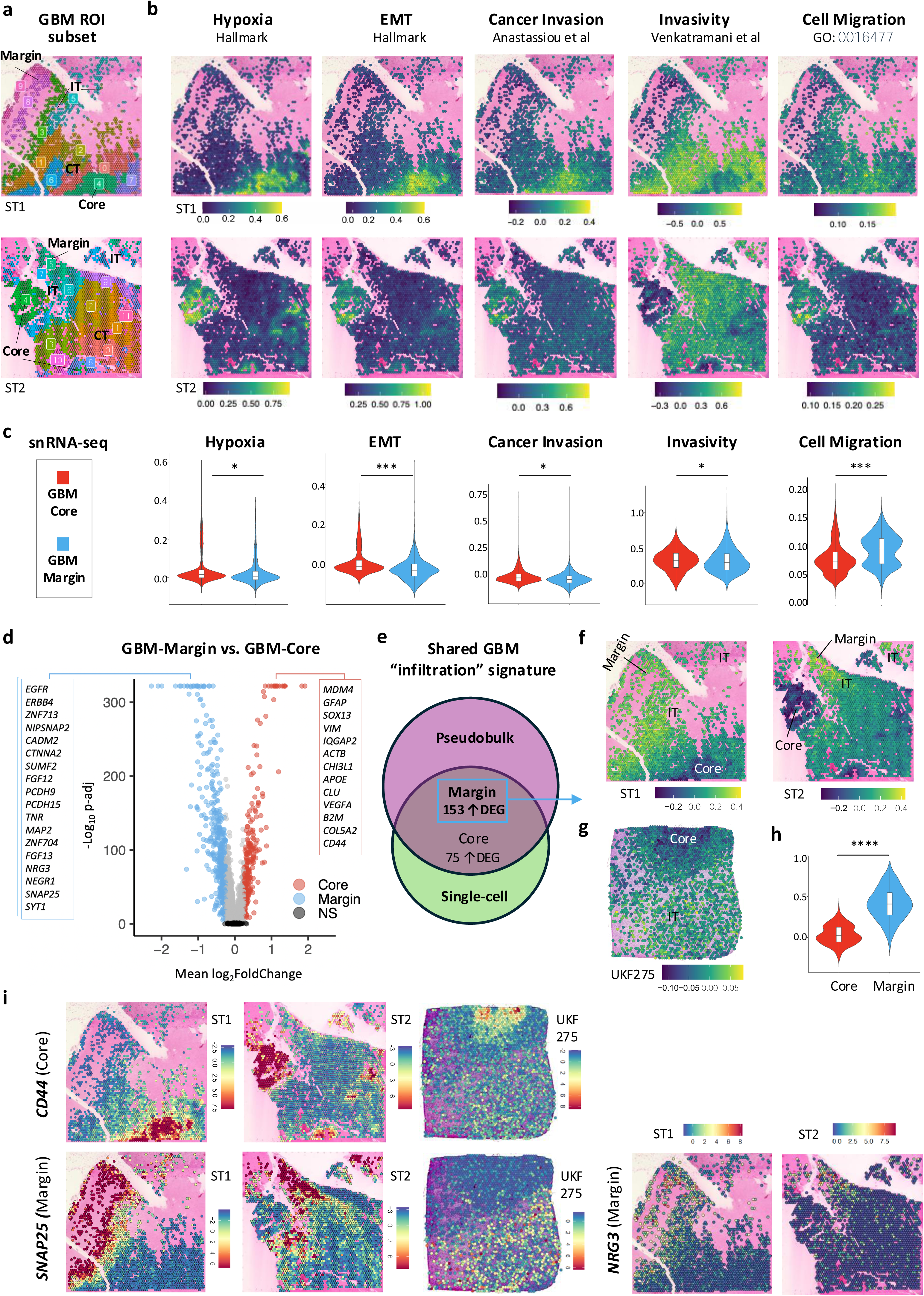
Derivation and spatial mapping of “GBM infiltration” signature. **a.** SpatialDimPlots of ST1 (top) and ST2 (bottom) subset for GBM-enriched ROI, colored by cluster number. **b.** SpatialFeaturePlots showing region-specific enrichment in ST1 (top) and ST2 (bottom) for pre-defined GBM signatures: “Hallmark hypoxia” in core; “Hallmark Epithelial Mesenchymal Transition (EMT)” in core; “Multicancer invasion” in core; “Invasivity” in CT and IT; and “GO:BP Cell migration” in CT and IT. **c.** Violin plots showing enrichment score of the same signatures in snRNA-seq GBM-Core and GBM-Margin populations, confidently dissected samples (* = p.adj < e^-100^, ** = p.adj < e^-200^, *** = p.adj < e^-300^, unpaired Wilcoxon rank test). **d.** Volcano plot showing differentially expressed genes (DEG) in GBM-Margin vs. GBM-Core analysis (p-adj < 0.05 and log2(FC) > 0.25). **e.** Venn diagram showing overlap of DEGs identified from single-cell level and pseudobulk-level GBM-Core vs. GBM-Margin differential expression analyses. **f.** SpatialFeaturePlot showing enrichment of “GBM infiltration” signature at the Margin in ST1 and ST2 samples. See also Supplementary Fig. 7c. **g.** SpatialFeaturePlot showing enrichment of “GBM infiltration” signature at IT zone in external ST sample UKF275 (Ravi et al). See also Supplementary Fig. 8. **h.** Violin plot showing significant enrichment of “GBM infiltration” in GBM-Margin populations from snRNA-seq data, confident dissected samples (**** = p.adj < e^-400^, unpaired Wilcoxon rank test). **i.** SpatialFeaturePlots of ST1 (left), ST2 (middle) and UKF275(right), validating expression of top DEGs identified in snRNA-seq analyses: *CD44* in core (top); *SNAP25* and *NRG3* in margin (bottom).

To define a shared GBM signature that specifically captures tumor cell biology at the distal resection margin, we leveraged our regionally-annotated GBM data and performed comprehensive differential gene expression analysis of GBM-Core vs. GBM-Margin populations, both at single-cell level and at pseudobulk level. Differential expression (DE) analysis at the single cell level, with subsampling and latent variable inclusion, identified 692 total differentially expressed genes (DEGs), with 420 DEGs upregulated in GBM-Margin (Fig. 3d, Extended Data 5). Pseudobulk level analysis defined 1001 DEGs, with 644 DEGs differentially upregulated in GBM-Margin (Extended Data 5). Among the upregulated genes in GBM-Margin populations were several related to OPC function (*PCDH9/15*, *TNR*, *LHFPL3*); neuronal function (*SYT1*, *SNAP25*, *MAP2*); cell-cell adhesion (*CDH10*, *NCAM2*, *ALCAM1*); glutamate receptor signaling (*GRIA2*, *GRID2*); and positive regulation of cell migration (*DOCK5*, *LGR6*; *FGF13*) (Fig. 3d, Supplementary Fig. 7b, Extended Data 5-7). Several cancer and development-related genes were also differentially enriched in GBM-Margin populations, including *EGFR*, *NRG3*, *ERBB4*, *PIK3R1*, *MAP3K1*, *CTNNA2/3, NFIB* and *RORB* (Fig. 3d, Extended Data 5-7). In contrast, genes upregulated in GBM-Core related to hypoxia, angiogenesis, apoptosis, gene expression, and inflammation (Fig. 3d, Supplementary Fig. 7b, Extended Data 8-9).

To increase specificity, we narrowed down our list of DEGs to those shared in both DE analyses: 153 upregulated DEGs in GBM-Margin and 75 upregulated DEGs in GBM-Core (Fig. 3e, Extended Data 10-11). We termed the 153 GBM-Margin DEG signature “GBM infiltration”, reasoning that these upregulated genes may reflect aspects of the tumor’s distal infiltrative biology. Projecting “GBM infiltration” onto ST1, ST2 and external ST datasets [31] showed stronger and preferential enrichment for distal IT and Margin GBM-enriched ROI (Fig. 3f-h, Supplementary Fig. 8), than seen with other known tumor invasion/migration signatures (Fig. 3a-c). GBM-Core signature was expectably enriched in the ST Core (Supplementary Fig. 7c, Supplementary Fig. 8).

Given the emerging contribution of the TME to therapeutic resistance and GBM progression [52-54], we also examined differential expression of non-neoplastic cell types between “Core” and “Margin” (Supplementary Fig. 7d-e). We found strong DEG Margin signatures for myeloid cells, BVCs, neurons, and oligodendrocytes (Supplementary Fig. 7d, Extended Data 12). Signature projections onto ST data confirmed spatial contributions of top candidates (Fig. 3i, Supplementary Fig. 7e), validating our data as a useful community resource for further investigation of tumor and TME biology at the Margin.

### Trajectory analysis of GBM transitioning from Core to Margin

We expected to find some constant and other dynamic changes in gene expression as tumor cells infiltrate brain parenchyma towards the margin of resection. To assess gene expression dynamics during GBM Core-to-Margin transition, we calculated tumor directionality using RNA velocity [55, 56] and constructed a pseudotime trajectory using monocle3 [57]. RNA velocity calculations based on ratios of unspliced to spliced RNA showed a predominant Core-to-Margin directionality (Fig. 4a, Supplementary Fig. 9a-b) and higher velocity length suggesting differentiation towards the Margin (Supplementary Fig. 9b). We then used monocle3 to order GBM nuclei along pseudotime, choosing “Core” as the starting point of transition (Fig. 4b, Supplementary Fig. 9b-c). To identify key genes involved in this transition, we analyzed differential RNA velocity expression and pseudotime progression modules from Core vs. Margin (Extended Data 13-15). There was significant enrichment of “GBM-infiltration” in top-ranked velocity and pseudotime Margin genes (Fig. 4c), prioritizing this subset of GBM “infiltration” genes as dynamically regulated during Core-to-Margin transition. Examples included *MAP2*, *TNR*, and *EGFR* (Fig. 4d, Extended Data 15).

**Figure 4:**
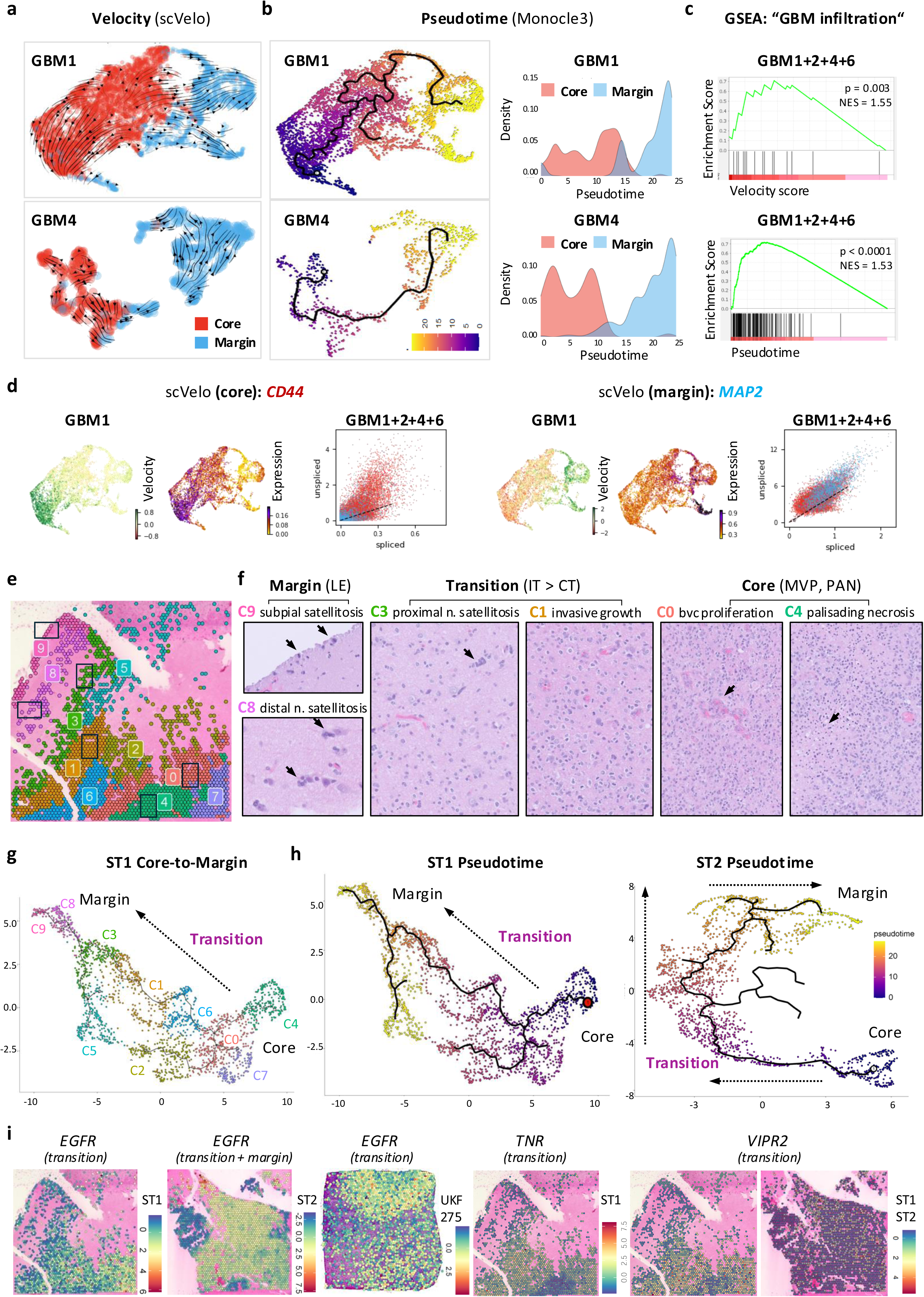
Inference of Core-to-Margin GBM infiltration trajectory and transition dynamics. **a.** Streamlines of RNA velocity on UMAP, calculated using the stochastic scVelo model, showing predominant core-to-margin directionality. Representative analysis for GBM1 (top) and GBM4 (bottom) are shown. See Supplementary Fig. 9a-b for other samples. **b.** UMAP (left) and density plot (right) of pseudotime, calculated using Monocle3, showing core-to-margin trajectory. Representative analyses for GBM1 (top) and GBM4 (bottom) are shown. See Supplementary Fig. 9b-c for other samples. **c.** Gene set enrichment analysis (GSEA) showing significant enrichment of “GBM infiltration” in a ranked list of “dynamic” genes ordered by RNA velocity (top) and pseudotime Moran’s test statistics (bottom) (GBM1+GBM2+GBM4+GBM6 combined). **d.** UMAPs of RNA velocity (left) and expression (middle) for top velocity score genes in GBM1 core (*CD44*) and margin (*MAP2*). Shown on right are the unspliced/spliced phase portraits for *CD44* and *MAP2*, across GBM-Core and GBM-Margin populations (GBM1+GBM2+GBM4+GBM6 combined). **e-f.** SpatialDimplot of ST1 GBM-enriched ROI with black rectangles (e) highlighting regions corresponding to specific histological features in (f), including subpial satellitosis, neuronal (n.) satellitosis, bvc proliferation, and palisading necrosis (marked by arrows). Clusters corresponding to each histology image are labelled with matching color codes. **g.** UMAP of ST1, with GBM-enriched ROI colored by cluster number. **h.** Pseudotime trajectory in ST1 (left) and ST2 (right). Arrows indicate direction of core-to-margin transitions. See also Supplementary Fig. 9d. **i.** SpatialFeaturePlots of GBM-enriched ROI for ST1, ST2 and UKF275, showing select genes (*EGFR*, *TNR*, *VIPR2*) identified in pseudotime-defined transition modules. See also Supplementary Fig. 9d-f.

While trajectory inference of snRNA-seq data informed the biology of GBM within its two opposing “Core” and “Margin” sectors, the analysis lacked representation of the intervening “transition” zone. We therefore used monocle3 in our spatial data to order GBM-enriched ROIs along a Core-to-Margin pseudotime trajectory, capturing clearly defined CT and IT populations within the intervening transition zone (Fig. 4e-h, Supplementary Fig. 9d). Differentially enriched genes within transition zone clusters, defined by monocle3 as modules (Supplementary Fig. 9d), included *EGFR*, *TNR*, and *VIPR2* (Fig. 4i) among others (Extended Data 16-17). Finally, we confirmed regional expression of transition and margin genes in external datasets, including at bulk level in IVY-GAP [58] (Supplementary Fig. 9e) and at single cell level [22] (Supplementary Fig. 9f).

Overall, trajectory analyses in our spatially resolved multi-omics dataset defined Core-to-Margin directionality and a subset of genes that are dynamically regulated in GBM cells during this transition. These represent potential targets in efforts to curtail the beginning of tumor infiltration, and understanding further their regulatory biology is critical.

### Distinct chromatin-level adaptations of GBM cells at the Margin

Changes in chromatin accessibility enable us to capture gene expression switches from poised to transcriptionally active states, and to infer TF activity that may underlie these changes [59]. To understand chromatin level changes that occur as infiltrative tumor cells adapt to the distinct “Margin” microenvironment, we turned to snATAC-seq performed in the same GBM-Core and GBM-Margin samples (Fig. 2b, Supplementary Fig. 2). We first analyzed differential chromatin accessibility in GBM-Core and GBM-Margin cells in relationship to their microenvironment (Extended Data 18-19). Broadly, we saw shared peak accessibility and associated TF motif activity between tumor and non-neoplastic populations, with distinctly co-opted states in GBM-Core vs. GBM-Margin (Fig. 5a, Supplementary Fig. 10a). Top TF motifs in GBM-Margin were shared with OPC and interneurons (such as ASCL1 and ASCL2); while those in GBM-Core were shared with BVCs and excitatory neurons (such as JUN and FOS members) (Fig. 5a). In contrast, both GBM-Core and GBM-Margin populations shared TF activity with astrocytes, including known astrocyte-driver NFIA [60] and RFX members (Fig. 5a).

**Figure 5:**
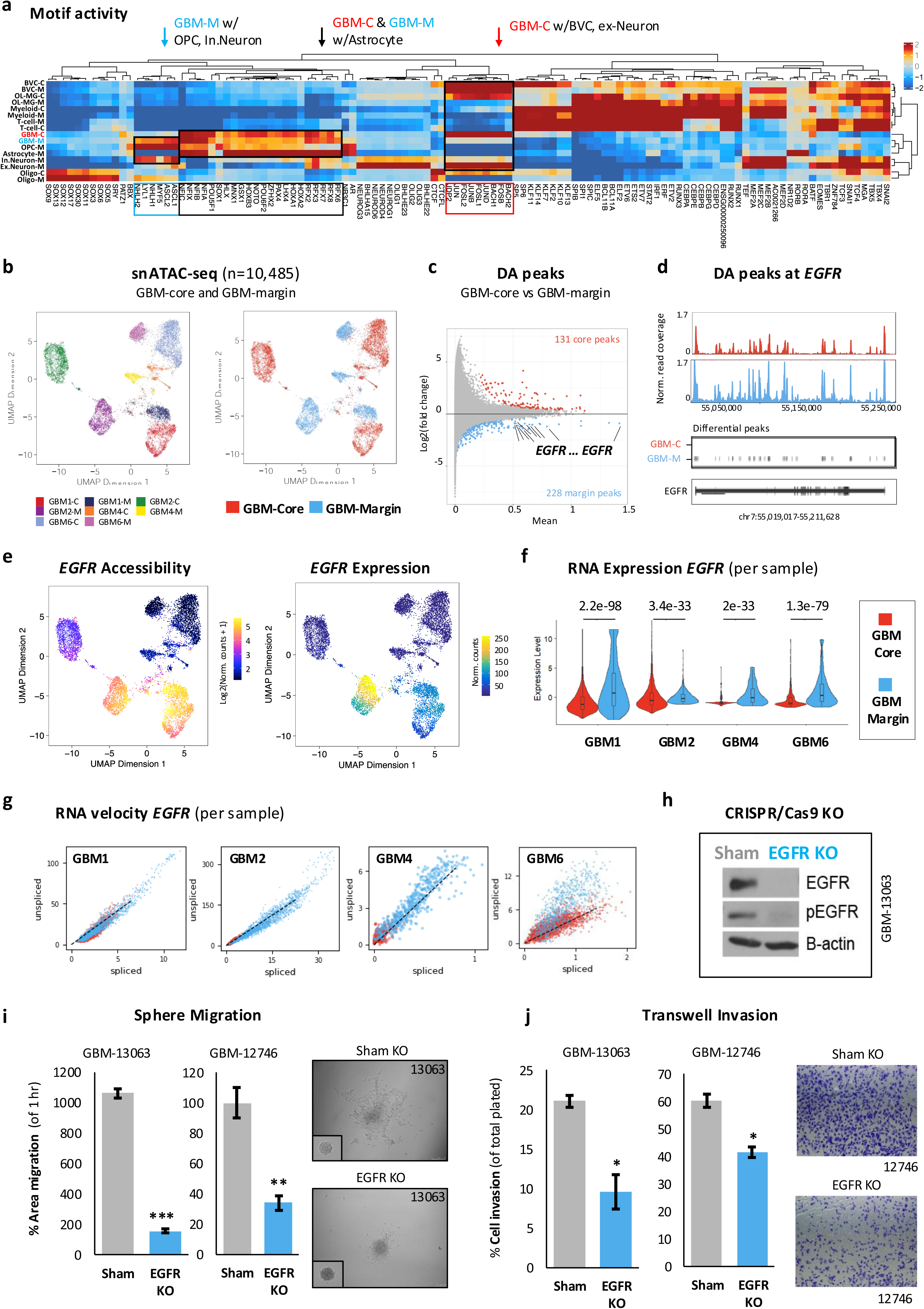
Prioritization of EGFR as top differential GBM-Margin marker and in-vitro validation of its role in GBM infiltration. **a.** Heatmap of top enriched motif activity from snATAC-seq data across all cell types in GBMs with confident core and margin dissections. (FDR < 0.05, MeanDiff >= 0.1, clusters with <100 cells omitted from analysis). See also Supplementary Fig.10a. **b.** UMAP of GBM-subset snATAC-seq data from a) colored by patient ID (left) and region (right). See also Supplementary Fig.10b. **c.** MA plot showing differential peaks identified in GBM-Core vs. GBM-Margin accessibility analysis (FDR < 0.05, abs(Log2FC) >= 0.1, abs(MeanDiff) > 0.2). Peaks mapping to EGFR locus are labelled. See also Supplementary Fig. 10c. **d.** Differential peaks at *EGFR* with corresponding read coverage in GBM-Core (top) and GBM-Margin (bottom). See also Supplementary Fig. 10e. **e.** UMAP showing corresponding *EGFR* accessibility and expression in GBM-subset snATAC-seq data. See also Supplementary Fig. 10f. **f.** Violin Plot of snRNA-seq data showing significantly higher *EGFR* expression in each GBM-Margin sample, compared to its GBM-Core (unpaired Wilcoxon rank test). **g.** Phase portraits of *EGFR* RNA velocity split by GBM sample and colored by region, showing higher velocity at each GBM-Margin sample, compared to its GBM-Core. **h.** Representative immunoblot images showing EGFR deletion and phospho-EGFR (pEGFR) loss after CRISPR/Cas9 knockout (EGFR KO) in patient-derived GBM-13063 cells, compared to Sham. ß-actin used as loading control. See also Supplementary Fig.10g. **i.** Bar plots (left) showing significantly diminished area of sphere migration at 36h in EGFRKO GBM-13063 and GBM-12746 cells, compared to Sham. Representative images (right) of spheres at 1h (inset) and 36h in Sham and EGFR KO G-13063. See also Supplementary Fig. 10h. **j.** Bar plots (left) showing significantly diminished percent transwell cell invasion in EGFRKO GBM-13063 and GBM-12746 GBM cells, compared to Sham. Representative images (right) of crystal violet-stained GBM-12746 cells invading across the transwell membrane, in sham (top) and EGFR KO (bottom). (* = p.val <0.05, ** =p.val < 0.01, *** =p.val < 0.001, unpaired two-tailed Student’s t-TEST for i and j)

To define tumor-specific differences in chromatin accessibility, we subclustered only GBM snATAC-seq data (Fig. 5b, Supplementary Fig. 10b) and performed differential accessibility (DA) peak analysis on GBM-Core vs. GBM-Margin (Fig. 5c). This defined a set of 131 GBM-Core and 228 GBM-Margin DA peaks (Fig. 5c, Extended Data 20) and TF motifs (Extended Data 21), with differential accessibility noted in several “GBM infiltration” genes. Interestingly, we observed robust differences in chromatin accessibility at large, chromosome-level regions between GBM-Core and GBM-Margin populations. Notably, top DA peaks in GBM-Margin cells localized to chromosome (chr) 7, while in GBM-Core cells - to chr1 (Supplementary Fig. 10c). This corresponded to CNV analysis from snRNA-seq data showing significantly higher chr1 gain in GBM-Core and significantly higher chr7 gain in GBM-Margin (Supplementary Fig. 10d).

### EGFR is a top differential GBM-Margin marker and a driver of tumor migration

The most frequent and top DA peak-associated gene in GBM-Margin populations was *EGFR*, encompassing the genomic region chr7:54,889,806-55,103,387 (Fig. 5c-d). Robust and differential accessibility at *EGFR* was most pronounced in samples GBM1 and GBM2, known to have amplified copies of the gene, but few DA peaks were also noted in GBM4-Margin, a tumor which did not carry *EGFR* amplification (Supplementary Fig. 10e, Extended Data 20). Importantly, *EGFR* expression and accessibility showed tumor specificity (Supplementary Fig. 1c, Supplementary Fig. 10f) and correspondence (Fig. 5e). Scrutinizing sample-dependent differences in our snRNA-seq data as well, we observed higher *EGFR* expression (Fig. 5f) and velocity (Fig. 5g) in each GBM-Margin sample, compared to its Core, in both EGFR-wildtype and EGFR-mutant cases, suggesting a shared role in GBM infiltration towards the margin.

EGFR is the most commonly mutated and overexpressed gene in GBM [1, 2, 61, 62], and has been widely studied for its role in gliomagenic growth and stem cell maintenance [63-65]. To validate the role of EGFR in GBM infiltration, we deleted EGFR using CRISPR/Cas9 genomic editing, in two primary patient-derived GBM cell lines, one of which carried EGFR copy number alterations (G-12746) and the other one which was EGFR-wildtype (G-13063) [37, 38] (Fig. 5h, Supplementary Fig. 10g). Knockout of EGFR (EGFR-KO) significantly reduced confluent GBM cell migration in sphere migration assays, compared to sham (Fig. 5i, Supplementary Fig. 10h). EGFR-KO cells also showed significantly lower invasion in transwell assays (Fig. 5j). To uncouple the biology of cell proliferation from that of cell infiltration in vitro, migration assays were performed in the absence of growth factors [38].

### The targetable regulator TEAD1 occupies accessible chromatin at *EGFR*

Studies from our lab and others have recently implicated the critical role of TEAD TFs and their co-activators YAP/TAZ in glioma stem cell plasticity [36], therapeutic resistance, and glioma migration and invasion [37, 38]. Given the implication of open chromatin poising *EGFR* expression in GBM-Margin cells, we leveraged our data to investigate further TEAD1’s binding activity at *EGFR*. Analysis of TEAD1 accessibility, motif activity, and predicted TF binding (footprinting) in snATAC-seq data confirmed tumor predominance, across both GBM-Core and GBM-Margin populations (Fig.6a-b, Supplementary Fig. 11a-b). Focusing specifically on GBM-Margin data where we find EGFR to be most accessible and expressed, snATAC-seq regulon analysis defined several potential TEAD1 cis-target genes, including *EGFR* (Fig. 6c, Supplementary Fig. 11c). Finally, to confirm trans-binding of TEAD1 at the EGFR locus, we performed ChIP-Seq with a specific TEAD1 antibody on all GBM specimens which showed accessibility at *EGFR*: GBM1, GBM2, and GBM4. Compared to IgG and input background controls, we observed strong enrichment for TEAD1 occupancy at *EGFR*, corresponding to open chromatin regions marked by H3K27ac and Tn5 accessibility (Fig. 6d, Supplementary Fig. 11d). TEAD1 occupancy signal was highest in EGFR-amplified GBM1 and GBM2 tumors, even after input-normalization (Fig. 6d, Supplementary Fig. 11d), corroborating findings from snATAC-seq data. Differential analysis of ChIP-Seq peaks showed significant enrichment at *EGFR* in GBM tumors compared to negative lymph node (LN) tissue control (Fig. 6e), which we confirmed by ChIP-qPCR (Supplementary Fig. 11e). Of note, we detected 5 differential TEAD1 occupancy peaks at the EGFR locus, more than for any other gene (Extended Data 22), prioritizing TEAD1 as potentially targetable regulator of EGFR expression.

**Figure 6:**
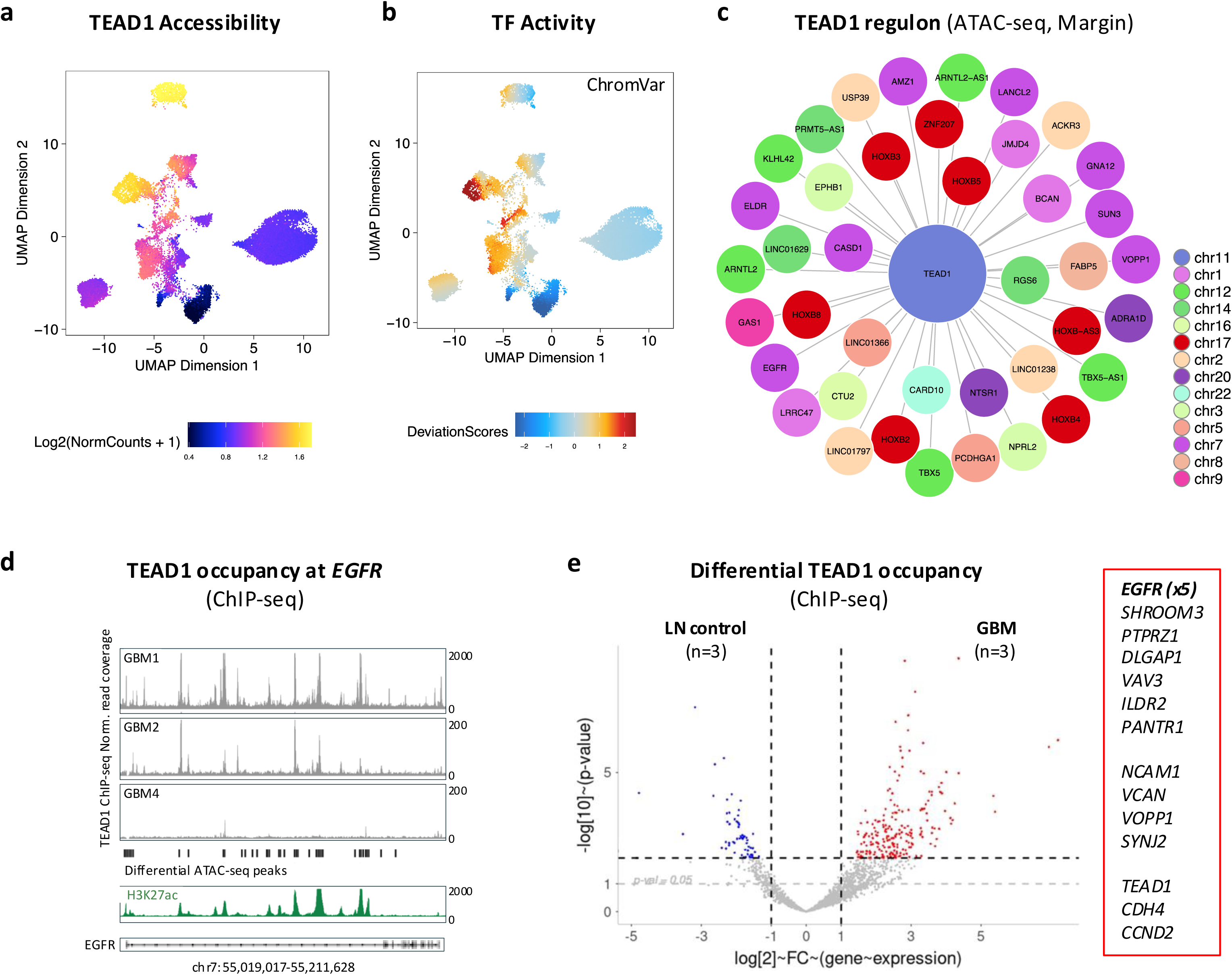
TEAD1 is an upstream regulator of EGFR. **a.** UMAP feature plot showing tumor-predominant TEAD1 accessibility. **b.** UMAP feature plot showing tumor-predominant ChromVar motif activity. See also Supplementary Fig. 11a-b. **c.** Regulon analysis of GBM-subset snATAC-seq data showing top TEAD1 target genes in GBM-Margin (correlation coefficient > 0.875). See also Supplementary Fig. 11c. **d.** TEAD1 occupancy analysis showing TEAD1 ChIP-seq read coverage at *EGFR* in GBM1, GBM2, and GBM4. Bottom panels show open chromatin defined by corresponding snATAC-seq differential peaks and H3K27ac ChiP-seq. See also Supplementary Fig. 11d. **e.** Volcano plot showing differential TEAD1 ChIP-seq peak occupancy, in negative control lymph node (LN) tissue vs GBM (n=3). Representative differential GBM TEAD1 peak-genes are shown in red box, some of which are also found in “GBM infiltration” signature. See also Supplementary Fig. 11e.

## Discussion

Most basic and preclinical studies have focused on GBM biology at the tumor core. Paradoxically, residual disease at the infiltrative margin predicts treatment response and drives recurrence [1, 6-8]. In this study, we used GBM tissue from rare supratotal neurosurgical resections [11] to generate a new multi-omic dataset that specifically profiles tumor and TME populations at the infiltrative “Margin” in addition to “Core”. To confidently distinguished normal from tumor populations at the margin, we implemented several technological improvements, including iterative cell-annotation efforts using external neurotypic cortex reference [46] and InferCNV [12], and subsetting GBM-enriched ROI in our ST-Visium analyses [42]. Combining ST data that spatially resolves the entire core-to-margin transition enabled us to confirm with high confidence our snRNA/ATAC dissections as well as to validate top differential markers. This multi-omic dataset builds on prior efforts characterizing invasive GBM biology at the rim and infiltrative edge [22-33], and provides the necessary extension to the most distal infiltrative biology in resected tissue that approximates the margin. Importantly, the dataset captures the full repertoire of TME cell types and enables the community to scrutinize further tumor and TME biology in the context of residual GBM disease.

To identify shared dependencies of infiltrative GBM cells despite tumor heterogeneity, we curated samples with diverse genomic drivers. By contrasting to Core, we define a robust, differential signature across snRNA-seq GBM populations at the infiltrative margin, termed “GBM infiltration”, and demonstrate its specific spatial enrichment near the margin, compared to prior gene sets that describe GBM invasion or migration [24, 51]. At the same time, we find similarities to prior GBM analyses focused on the “periphery” or “rim” [23, 24, 27, 29], including OPC-like and neuronal-like enrichment, consistent with infiltrative biology representing a continuum rather than a unique on/off state [28, 30, 31]. Previous studies have suggested that GBM infiltration biology is sustained by adaptive plasticity and is maintained at the level of open chromatin [10, 66]. Consistent with that, we reconstructed a core-to-margin trajectory confirming directionality and differentiation towards the margin, and then defined dynamically regulated markers during this transition as well as a unique accessibility signature in GBM-margin populations. We found EGFR to be the top tumor-specific marker in these analyses, across both EGFR-wildtype and EGFR-mutant tumors. Accessibility was noticeably higher in GBM with EGFR amplification, correlated to but exceeding inferred chr7 gain. This is consistent with prior studies demonstrating unique accessibility in GBM subpopulations with focal amplification [67] and is at least partially explained by the rudimentary topology of open chromatin at extrachromosomal DNA (ecDNA) [68-70]. Importantly, it suggests that chromatin accessibility at EGFR ecDNA may be sustaining infiltrative GBM biology at the margin, where clonal selection of EGFR-amplified GBM cells has been previously observed [71, 72]. The importance of EGFR in stem cell maintenance, proliferation, and gliogenesis is well established [46], with few studies also suggesting a role in glioma invasion [73-76]. Here we deleted EGFR using CRISPR/Cas9 to also validate EGFR’s role in GBM migration using patient-derived models.

While EGFR is silenced in adult brain, it is aberrantly re-expressed in most high-grade gliomas, even in those without EGFR amplification, through a mechanism that includes aberrant chromatin remodeling at its promoter [62]. Defining upstream regulators of *EGFR* transcription may thus provide new therapeutic opportunities, especially since adaptive resistance of GBM cells to downstream signaling has rendered traditional tyrosine kinase inhibitors ineffective [77]. Here we prioritized TEAD1 as tumor-predominant TF with EGFR cis-activity in snRNA/ATAC-seq data, and then confirmed its trans-binding activity through genome-wide ChIP-seq analysis. TEAD1 occupancy at EGFR was noted across GBMs, with the signal being strongest in EGFR-amplified tumors, even after input normalization, consistent with ATAC-seq findings. TEAD1 and its co-activators YAP/TAZ are critical drivers of GSC plasticity [36] and migration/invasion [37, 38], with emerging evidence for crosstalk between Hippo YAP-TEAD and EGFR emerging [37, 78-81]. Given the excitement of YAP-TEAD inhibitors as new GBM therapeutics in preclinical and early clinical trial studies [82], their therapeutic efficacy at the resection margin deserves important consideration in future studies.

## Methods

### Sample collection

Sample collection was conducted in a de-identified manner, under appropriate consent, and in compliance with the policies and regulations at the Icahn School of Medicine at Mount Sinai and its institutional review board. Surgical glioma tissue was collected from 8 patients with confirmed diagnosis of glioblastoma, WHO grade 4 (2016 and 2021 diagnostic criteria), representing diverse genetic backgrounds and all major mutational drivers. Tissue procurement was coordinated with neurosurgery to include contrast-enhancing tumor “Core” and FLAIR-hyperintense infiltrative “Margin” representing the distal extent of gross supratotal resection and having a “normal brain” appearance on fresh tissue inspection. Experienced neuropathologist confirmed the presence of necrosis in “Core” and the presence of infiltrative brain parenchyma without necrosis in “Margin” tissues, during frozen section dissection. Sections were snap-frozen, stored at -80°C for snRNA-seq (n=6) and/or fixed in 10% formalin for spatial transcriptomics (n=2). Serial formalin-fixed and paraffin embedded (FFPE) sections from samples were kept for histological analysis.

### Nuclei isolation

Nuclei were isolated separately from each GBM “Core” and “Margin” fresh-frozen tissue dissection, and the same dissociate preparation was used for downstream snRNA-seq and snATAC-seq library preparations using published protocols [46]. Briefly, 50-200 mg of tissue was dounced in 4 ml of lysis buffer (0.32 M sucrose, 5 mM calcium chloride, 0.1% Triton X-100, 0.1 mM ethylenediaminetetraacetic acid [EDTA], 3 mM magnesium acetate, and 1 mM dithiothreitol [DTT] in 1 mM Tris-HCl, pH 8.0) under RNase-free working conditions. Nuclei were isolated by sucrose gradient ultracentrifugation: a sucrose solution (1.8 M sucrose, 3 mM magnesium acetate, and 1 mM DTT in 10 mM Tris-HCl, pH 8.0) was layered under the lysate solution followed by centrifugation at 4°C for 1 hour (h) at 24,400 rotations per minute. Nuclei were resuspended in 0.04% bovine serum albumin (BSA) in Ca2+/Mg2+-free phosphate buffered saline (PBS) at 1x10^6^ cells/mL concentration and loaded to achieve optimal recovery of 6,000 sequenced nuclei/sample with minimal doublet/triplet formation. Concentrations were verified using an automated cell counter (Countless 3, ThermoFisher). Trypan blue staining was used to assess nuclei quality and quantity prior to library preparation.

### Library preparation and high-throughput sequencing

#### snRNA-seq

RNA from single nuclei was prepared for sequencing using the 10X Genomics Chromium platform and the 3’ gene expression (3’ GEX) kit, v3.10. Each region (Core and Margin) and sample were barcoded and sequenced as separate libraries, without hashing. Nuclei were diluted and loaded with a target of 6,000 cells into nanoliter-scale Gel Bead-In-Emulsions (GEMs). Primers containing Illumina read 1 (R1) sequencing primers, a 16-bp 10x Barcode, a 10-bp randomer and a poly-dT primer sequence were subsequently mixed with the nuclear suspension and master mix. After incubation of the GEMs, barcoded, full-length cDNA was generated from pre-mRNA. Then, the GEMs were broken, and silane magnetic beads were used to remove leftover biochemical reagents and primers. Prior to library construction, enzymatic fragmentation and size selection were used to optimize cDNA amplicon size. P5 and P7 primers, a sample index, and Illumina read 2 (R2) sequencing primers were added to each selected cDNA during end repair and adaptor ligation. P5 and P7 primers were used for Illumina bridge cDNA amplification (http://10xgenomics.com). Libraries were quantified using the Agilent Bioanalyzer and sequenced into 2x100 paired end reads using the Illumina NovaSeq platform.

#### snATAC-seq

Nuclei were transposed and processed according to Chromium Single Cell ATAC kit (10X Genomics, Pleasanton, CA) using manufacturer’s guidelines. An estimated 10,000 nuclei were loaded onto each channel with a targeted nuclei recovery of 6000 nuclei/sample. Fragmented DNA underwent RT and cleanup followed by amplification 10X-specific sample indexing following the manufacturer’s protocol. Libraries were quantified using Tapestation (Agilent) and Qubit (ThermoFisher) analysis. Libraries were sequenced in paired end mode on a NovaSeq instrument (Illumina, San Diego, CA) targeting a depth of 50,000-100,000 reads per cell.

#### Spatial Transcriptomics (ST)

10X-Visium was used to perform ST on two GBM specimens, with distinct genetic background and patient sex, in which highest RNA quality was obtained and full Core-to-Margin transition was captured histologically. Tissue permeabilization and library preparation was done according to the manufacturer’s protocol. The Visium spatial gene expression for FFPE assays (10X Genomics) was used to generates spatially-resolved cDNA libraries from FFPE tissue. Tissue slices were sectioned onto the assay slide, which consists of four 6.5mm x 6.5mm arrays of spatially-barcoded poly T capture probes (arranged as 5,000 55 um spots) and permeabilized. Captured mRNA then underwent reverse transcription and cleavage from the slide, followed by cDNA amplification and fragmenting, end-repair, poly A-tailing, adapter ligation, and 10X-specific sample indexing following the manufacturer’s protocol. Libraries were quantified using Tapestation and Qubit and sequenced in paired end mode on NovaSeq instrument targeting a depth of 50,000-100,000 reads per capture spot.

### Sequence alignment and quality control

10X Genomics Cell Ranger (v.7.1.0) and Space Ranger (v2.1) were used respectively for snRNA-seq/snATAC-seq and ST data demultiplexing, processing, and GRCh38 (hg38) genome assembly along with Ensemble transcript annotations. Default parameters were used unless specified. For both snRNA-seq and ST, feature-barcode matrices were processed, and quality-controlled using Seurat package (v4.3.0) [41, 42]. Low quality nuclei were removed based on three different criteria: a) nuclei with percent mitochondrial gene reads greater than 15%, b) nuclei with less than 400 features, and c) nuclei with fewer than 1000 counts. We considered the fact that GBM cells might have higher mitochondrial activity than normal cells and that neurons express many more genes than other cell types when applying this criterion. To remove potential doublets from snRNA-seq and ST data, we filtered out nuclei with greater than 10,000 features and additionally used *Doubletfinder* (v3 with *paramSweep_v3*). For snATAC-seq, fragment files were processed using the ArchR package (v1.0.1) [47]. Nuclei were filtered based on TSS enrichment less than 4 and fragment number less than 3000 and greater than 30,000. Doublets were predicted and removed using *addDoubletSCores* and *filterDoublets*, respectively.

### Multi-omics data normalization and integration

#### snRNA-seq

After QC filtering, snRNA-seq per sample counts were normalized and scaled using *SCTransform* function setting *vst.flavor* to v2, which employs the *glmGamPoi_offset* method. This process included subsampling with 2000 cells and excluding Poisson genes from learning and regularization. *SelectIntegrationFeatures* was used to select 3000 top scoring features across datasets. *PrepSCTIntegration* was used to ensure the features selected were present in the objects-list used and *FindIntegrationAnchors* was used to set anchors between objects in the list, followed by integration of objects using *IntegrateData* function. Dimensionality reduction was performed using principal component analysis (PCA), followed by Uniform Manifold Approximation and Projection (UMAP) embedding for visualization. Nearest neighbors were computed with default parameters and clusters identified using Seurat’s *FindClusters* function, which employs shared nearest neighbor modularity optimization.

#### ST

Output from 10X genomics Space Ranger pipeline was imported into Seurat using *Load10X_Spatial* to generate a separate Seurat object for each ST. Dimensionality reduction using PCA, clustering and UMAP embeddings were generated using default parameters as above, including SCTransform for normalization.

#### snATAC-seq

Dimensionality reduction was performed using the default *TileMatrix* and the *addIterativeLSI* function with 15,000 variable features, 2:30 dimensions, 10000 sample cells, 15 maximum clusters, n.start 10, and 0.5 cluster resolution. Clusters were computed using *addClusters*. Dimensionality reduction and clustering were repeated with the *PeakMatrix* using the same parameters described above. Batch effect correction was performed using Harmony [48] followed by *addUMAP* and *addClusters*.

### Annotation of cell type and state

#### snRNA-seq

Cell-type annotation was based on canonical markers for normal brain cell types and glioma stem cells (GSC), calculated from top marker gene expression averaged per cluster, as well as by projection of snRNA-seq neurotypic adult cortex [46] cell type signatures. InferCNV [12] with default settings was used to infer copy number variation (CNV) status in tumor populations from snRNA-seq data, including gain of chromosome 7 (+7) and loss of chromosome 10 (-10), using as references both internal normal cells (neurons + myeloid) and external neurotypic cortex dataset [46]. A combined score including +7/-10 CNV and GSC markers (“tumor status”) was used to further increase the specificity of malignant cell annotations. Cell cycle scoring was performed in Seurat using *CellCycleScoring* and established cell cycle phase markers to classify nuclei into G2/M, S and G1 phase [83]. Cell type or cluster proportions were calculated as a ratio of the total number of nuclei per region.

#### snATAC-seq

Clusters were annotated by cell type using marker gene accessibility (GeneScoreMatrix) and expression (GeneIntegrationMatrix) after computing imputations weights with *addImputeWeights*. Pseudo-bulk replicates were created by cell type using *addGroupCoverages* and a reproducible peak set was generated with *addReproduciblePeakSet*, calling peaks with MACS2 (v2.2.9.1) [84]. Integration with snRNA-seq data from corresponding samples was performed with the *addGeneIntegrationMatrix* function. Peak-to-gene links were computed using the *addPeak2GeneLinks* function and heatmaps were plotted with *plotPeak2GeneHeatmap* (k = 15). CNVs were predicted using snRNA-seq label transfer by matching corresponding nuclei by barcode.

#### ST

For tumor CNV prediction in spatial data, the InferCNV pipeline in SPATA2 v2.0.4 (*runCNVanalysis* function) was used. For cell type deconvolution, RCTD [49] and *FindTransferAnchors* in Seurat [42] were used to assign a unique cell type to each ROI and predict GBM-enriched ROI spots.

### Differential gene, peak, and motif analyses

#### snRNA-seq

Differential gene expression analysis (DGEA) was performed at both single-cell and pseudobulk levels. For single-cell level DGEA, we used Seurat’s *FindMarkers* function, with logistic regression method and specifying ‘patient’ as the latent variable for the test. Additionally, we randomly subsampled cells, taking the lowest number of cells in conditions that were compared, performed 1000 iterations using default parameters, and shortlisted genes that appeared more than 900 times with p-adj. < 0.05. For pseudobulk level analysis, we shortlisted differential genes based on p-adj. < 0.05 and abs(Log2FC) >= 0.5.

#### snATAC-seq

Differential analyses were performed using *GetMarkerFeatures*, using the Wilcoxon test, TSS enrichment and log10Fragments as bias, and default max cells value of 500. Significant marker genes were selected using *getMarkers* with the following cut-off values: FDR <= 0.01 & Log2FC >= 0.5 for GeneScoreMatrix and GeneExpression Matrix markers; FDR < 0.05, abs(Log2FC) >= 0.1, abs(MeanDiff) > 0.2 for differential peaks; and FDR < 0.05, MeanDiff >= 0.1 for differential motifs. Motif annotations from the CIS-BP database [85] were added using *addMotifAnnotations* and a deviation matrix was calculated with the *addDeviationsMatrix* function, using chromVAR in R (v1.18.0) [86]. Footprints were retrieved using *getPositions* and *getFootprints* and plotted with *plotFootprints*. TEAD1 target genes were identified by extracting peak coordinates from *getMatches* in CIS-BP and selecting for peaks with a peak-to-gene correlation coefficient > 0.87. Cell groups with fewer than 100 cells were excluded from differential analyses.

### Gene expression enrichment analyses

The *AddModulesCore* function in Seurat was used to calculate average gene expression / expression enrichment scores for single genes or gene sets, which were then projected onto snRNA-seq GBM-Core/Margin and onto ST Core-to-Margin data for spatial context. Gene sets included pre-defined Hallmark “hypoxia”, Hallmark “angiogenesis”, Hallmark “EMT”, Gene ontology (GO) “cell migration”, “multicancer invasion” [51], GBM “invasivity” [24] and herein defined “GBM infiltration”. GBM “invasivity” was defined as the top 20 differentially expressed genes in unconnected GBM populations [24] and “GBM infiltration” was defined as the 153 differentially expressed genes in GBM-Margin (vs. GBM-Core), shared between pseudobulk and single-cell snRNA-seq DGEA. An unpaired Wilcoxon rank test was used to calculate statistical differences between module scores in GBM-Margin vs. GBM-Core.

To assess molecular heterogeneity in our snRNA-seq data clusters, we performed single cell gene set enrichment analysis (ssGSEA) using Escape v2. Hallmark gene sets used were imported from Molecular Signature Database (MsigDB) using *getGeneSets* function in Escape. Enrichment score matrix was generated from SCTransformed count data for individual cells for selected gene sets using *escape.matrix* function and ssGSEA method along with normalization set to TRUE. EnrichR [87] was used for GO term enrichment analysis of differentially expressed genes in GBM-Core vs. GBM-Margin.

### External dataset analysis

For all external data enrichment analyses, a curated list of genes from published data [16, 18, 24, 28, 51] was used along with the *AddModuleScore* function to calculate enrichment scores. Two-dimensional representation of Neftel et al [16] data was generated using enrichment scores calculated for each cell state and projected onto each nuclei, using ggplot2. Briefly, the values on the Y axis represent the maximum score from OPC-like and AC-like states subtracted from NPC-like and MES-like states. Values on X-axis represent OPC-like minus maximum of NPC-like score for Y-axis greater than 0 and for Y-axis less than 0, the X-axis values represent AC-like minus maximum of MES-like score.

### Trajectory inference using RNA velocity and pseudotime

#### RNA velocity

Sample-specific loom files were generated from Cell Ranger output files using the Python package, *velocyto* (v0.17.17) [56]. RNA velocity and trajectory direction were inferred using the Python package *scVelo* (v0.2.5) [55]. Anndata (v0.8.0) was used to concatenate loom files. These were merged with Anndata objects created using the embeddings and count data of processed Seurat objects. Filtering was based on top 2000 variable features and 20 minimum shared counts, followed by normalization. Velocities were calculated using a stochastic model, and projected as streamlines or features onto a pre-computed UMAP embedding. Expression Velocity scores were used to rank the top 100 genes dynamically regulated in Core versus the Margin.

#### Pseudotime

Monocle3’s *learn_graph* function was used to define a trajectory path in snRNA-seq and ST data, and the *order_cells* function was used to order the cells along this path. We chose GBM-Core enriched cluster as the root node. The function *graph_test*, with the neighbor graph parameter set to *principal_graph*, was used to correlate region (Core or Margin) with gene expression in pseudotime. Genes filtered by q_value < 0.05 were selected from the results of graph*_test* to identify modules, composed of co-regulated genes. We used the *find_gene_modules* function to identify modules, choosing the lowest resolution that clearly distinguished differential co-regulatory modules in Core, Transition, and Margin.

### Data handling and visualization

Data was handled using Python (v 3.7.12) and R (v4.1.0). Associated packages included SciPy (v1.7.3), Scanpy (v1.9.3), NumPy (v1.21.6), and Pandas (v1.3.5) for Python and dplyr (v1.1.4) and collapse (v2.0.10) for R. Data were visualized using the Python package Matplotlib (v3.5.3), the R packages ggplot2 (v3.5.0), ggrepel (v0.9.1), ComplexHeatmap (v2.14.0), igraph (v1.4.1), EnhancedVolcano (v1.14.0), and built-in Seurat and ArchR functions such as *VinPLot* for violin plot visualization.

### TEAD1 occupancy analysis

TEAD1 occupancy was assessed via chromatin immunoprecipitation followed by genome-wide sequencing (ChIP-seq), using ChIP-grade TEAD1-specific antibody (BD Biosciences 610923, 5 μg) [37, 88]. TEAD1-ChiP-seq was performed on fresh-frozen patient-derived GBM tissue, including from GBM1, GBM2, and GBM4 also used for multi-omics, as well as on lymph node (LN) tissue with minimal TEAD1 expression to serve as negative control (n=3 technical replicates). ChIP-seq for open chromatin histone marker H3K27ac (Abcam ab4729, 5 μg) was also performed in a subset of samples. IgG ChIP-seq (12–371, Millipore, 5 μg) was performed as non-specific binding control, and 10% input was sequenced for DNA copy number normalization. ChIP assays were performed as previously described [37, 62]. Briefly, up to 400mg of GBM or LN tissue was thawed, minced, briefly cross-linked in 1% formaldehyde for 10 minutes, quenched in 2M Glycine for 5 minutes, lysed in 1X PBS containing PMSF and protease inhibitor cocktails (1%SDS, 50mM Tris-HCl, ph-8.0, 10mM EDTA, 1X PIC, 1mM PMSF) on ice for 15 min, and then homogenized using a Dounce homogenizer. Chromatin was fragmented to 300-500bp using sonication. Specifically, lysed samples were distributed into Eppendorf tubes to a maximum of 150 μl per tube and sonicated using a Bioruptor (30” ON/OFF per cycle – 50 cycles for GBM samples and 70 cycles for lymph node tissue). Following sonication, the samples were centrifuged at 12,000g for 10 minutes and the final lysate was carefully collected in a new tube. 10% of each lysate was reverse crosslinked and the extracted DNA was run on a 1.5% agarose gel to confirm fragment size. Dynabeads (35 μl of anti-Mouse or anti-Rabbit for TEAD1 and H3K27ac antibodies, respectively) per ChIP were blocked in 0.5% BSA, washed 3x in RIPA buffer and resuspended in blocking solution (100 μl per ChIP). 5 μg of antibodies for H3K27ac, TEAD1 and negative control IgG were added to the respective tubes followed by incubation for 6h at 4°C on a rotator. 20,000-25,000 ng of lysate was used per ChIP. Antibodies bound to magnetic Dynabeads were washed and resuspended in RIPA buffer followed by addition to the respective lysate-containing tubes. The reactions were incubated for 14h on a rotator at 4°C. After incubation, the reactions were washed with RIPA buffer and TE Buffer, and reverse crosslinked on a thermomixer for 3h using elution buffer and Proteinase K. ChIP DNA was extracted using QiaQuick PCR Purification kit (Qiagen) and quantified using Qubit Fluorometer 4.0.

ChIP-seq libraries were generated using the NEBNext® Ultra™ II DNA Library Prep Kit for Illumina® (NEB #E7645S). Size selection was carried out in samples with > 50 ng of total DNA. PCR enrichment of adaptor-ligated DNA was carried out using NEBNext Multiplex Oligos for Illumina #E7335 and #E7500 and the number of cycles was dependent on total amount of DNA at the start of library preparation. The libraries were run on a Bioanalyzer using the Agilent High Sensitivity DNA Kit to determine fragment size distribution. The libraries were sequenced at the Genomics Core Facility (Icahn School of Medicine at Mount Sinai) using the NovaSeq 6000 system as 50 bp paired end reads at a depth of ∼50 million reads (ChIP) and 25 million reads (input). Peaks per sample were called using MACS2, filtered for duplicates and blacklisted regions, and normalized per sample to their respective genomic input. Integrative Genomics Viewer (IGV) was used to visualize peaks. Quantitative *PCR (*qPCR) was performed on the post-ChIP DNA for validation using PerfeCTa® SYBR® Green FastMix®, ROX™ and previously published primers [37]. Enrichment was calculated as fold increase over IgG, after normalization with 10% input, using 2^–ΔΔCt^ analysis. The average Ct value from two replicates was used for the analysis.

### EGFR deletion and in-vitro functional validation studies

EGFR was deleted using CRISPR/Cas9 editing by introducing stop codon mutation in exon1, as previously described [37]. Two different gRNAs targeting exon1 were used generate population knockout (EGFR-KO) (F1 5’-CACCGTCCTCCAGAGCCCGACTCGC-3’ and R1 5’-AAACGCGAGTCGGGCTCTGGAGGAC-3’; F2 5’-CACCGGCGACCCTCCGGGACGGCCG-3’ and R2: 5’-AAACCGGCCGTCCCGGAGGGTCGCC-3’). For negative control, sham CRISPR/Cas9 knockout was generated targeting the non-human GFP gene (Sham-KO) (F: 5’-CACCGGGGCGAGGAGCTGTTCACCG-3’ and R: 5’-AAACCGGTGAACAGCTCCTCGCCCC-3’).

All in vitro validation experiments were performed in low-passage primary GBM cell lines (GSC-like), grown on laminin (10 μg/ml) in serum-free neutrosphere DMEM media supplemented with EGF (20 ng/ml) and bFGF (20 ng/ml). To form spheres, patient-derived GBM cells (sham-KO and EGFR-KO) were cultured on 96-well ultra-low adherence plates at a density of 2.5 cells/μl. Sphere diameter and number were recorded 6 days later.

Infiltration was evaluated using spheroid migration and transwell invasion assays. For spheroid migration, spheres were placed on poly-D-lysine (PDL) (10 μg/ml) with or without laminin substrate in serum-free conditions, and without EGF or bFGF growth factors. Spheroid migration was assessed at baseline (1h) and at 36h, by measuring sphere diameter and the area of confluent cell migration. For transwell invasion assays, approximately 25,000 live cells were plated in the top chamber of the transwell (Corning, 3470) without growth factors (containing only DMEM/F12, 0.6% D-Glucose, 2 mM L-Glutamine, 1× Antibiotic–Antimycotic, 15 mM HEPES) and allowed to invade through a laminin-coated (20 μg/ml) porous membrane. Growth factor and serum attractants were placed in the bottom chamber (DMEM/F12, 1× N2, 1× B27, 1× ITS, 0.6% D-Glucose, 2 mM L-Glutamine, 1× Antibiotic–Antimycotic, 15 mM HEPES, EGF (20 ng/ml), bFGF (20 ng/ml), 10% FBS). Percent cell invasion was measured at 24h by quantifying the number of invasive cells (adherent to the bottom of the membrane) out of the total cells plated.

---

**Supplementary Fig. 1:**
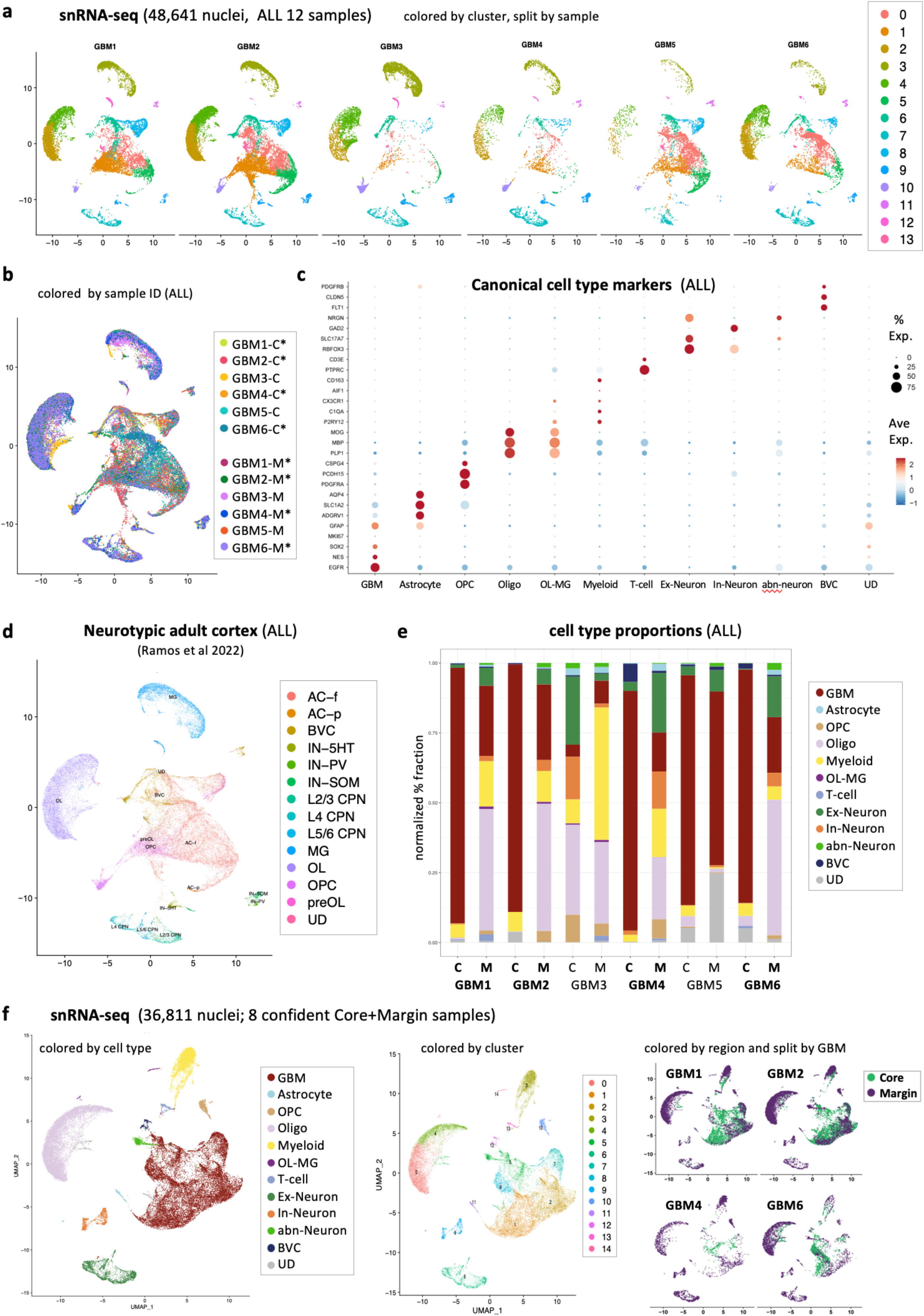
(relates to Fig. 1 and Fig. 2) **a.** UMAP of snRNA-seq data from all GBM samples (n=6), integrated, colored by cluster number and split by patient ID. **b.** UMAP of snRNA-seq data from all GBM samples (n=6), integrated, colored by patient ID and region (C=Core; M=Margin). **c.** Dot plot of canonical markers used for cell-type annotation, snRNA-seq data (n=6). **d.** UMAP of snRNA-seq data from all GBM samples (n=6) showing predicted cell types using adult neurotypic cortex as a reference. **e.** Stacked bar plot showing cell type proportions calculated for each snRNA-seq data sample, in Core and in Margin. **f.** UMAP of snRNA-seq data from confidently dissected Core and Margin GBM samples (n=4), integrated, colored by cell type (left), by cluster number (middle) and by region and split by patient ID (right).

**Supplementary Fig. 2:**
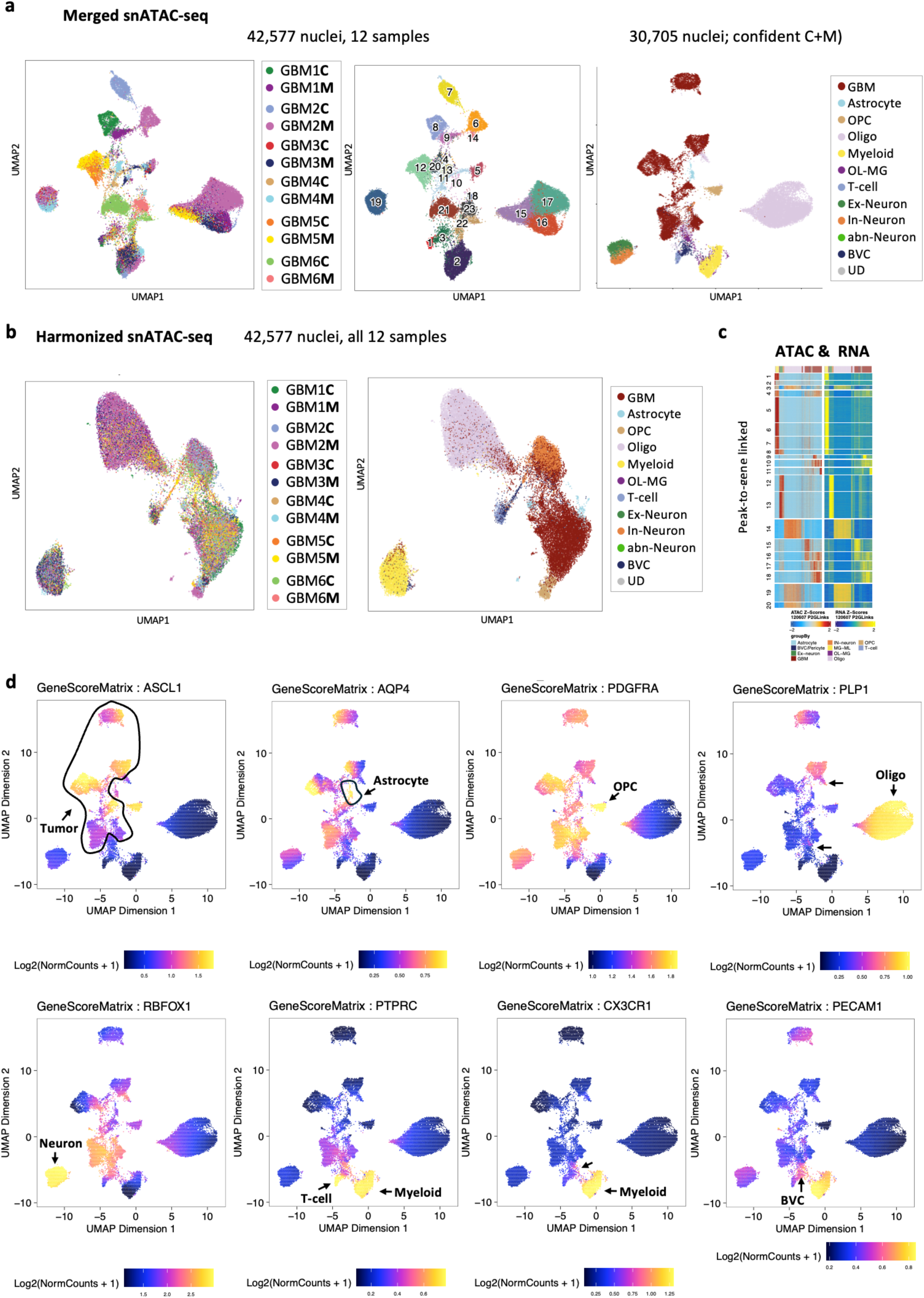
(relates to Fig. 1 and Fig. 2) **a.** UMAP of merged snATAC-seq data from GBM samples, colored by patient ID and region (left), cluster number (middle), and by cell type (right). **b.** UMAP of harmonized snATAC-seq data from all GBM samples (n=6), colored by patient ID and region (left) and by cell type (right). **c.** Heatmap of peak-to-gene linked snATAC-seq data showing cross-platform integration (confident core and margin GBM samples shown). **d.** Feature plot UMAPs of top marker gene scores used for cell type annotations, in merged snATAC-seq data from confidently dissected core and margin GBM samples (n=4).

**Supplementary Fig. 3.**
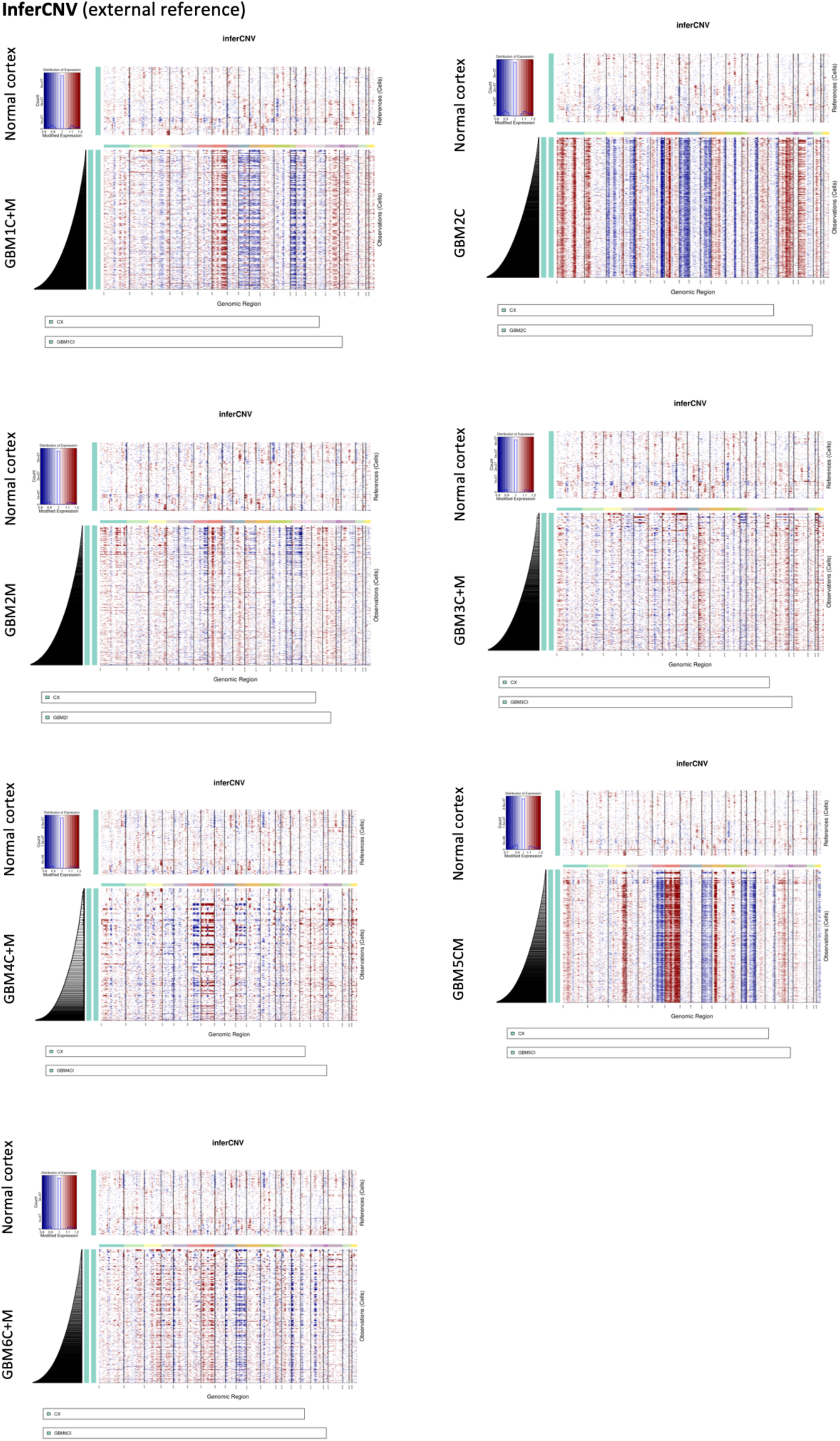
(relates to Fig. 1) Heatmaps showing inferred copy number variations using InferCNV for each GBM sample using external normal cortex dataset as a reference. For GBM2, heatmaps are split by patient ID and by region. C= core, M = margin.

**Supplementary Fig. 4:**
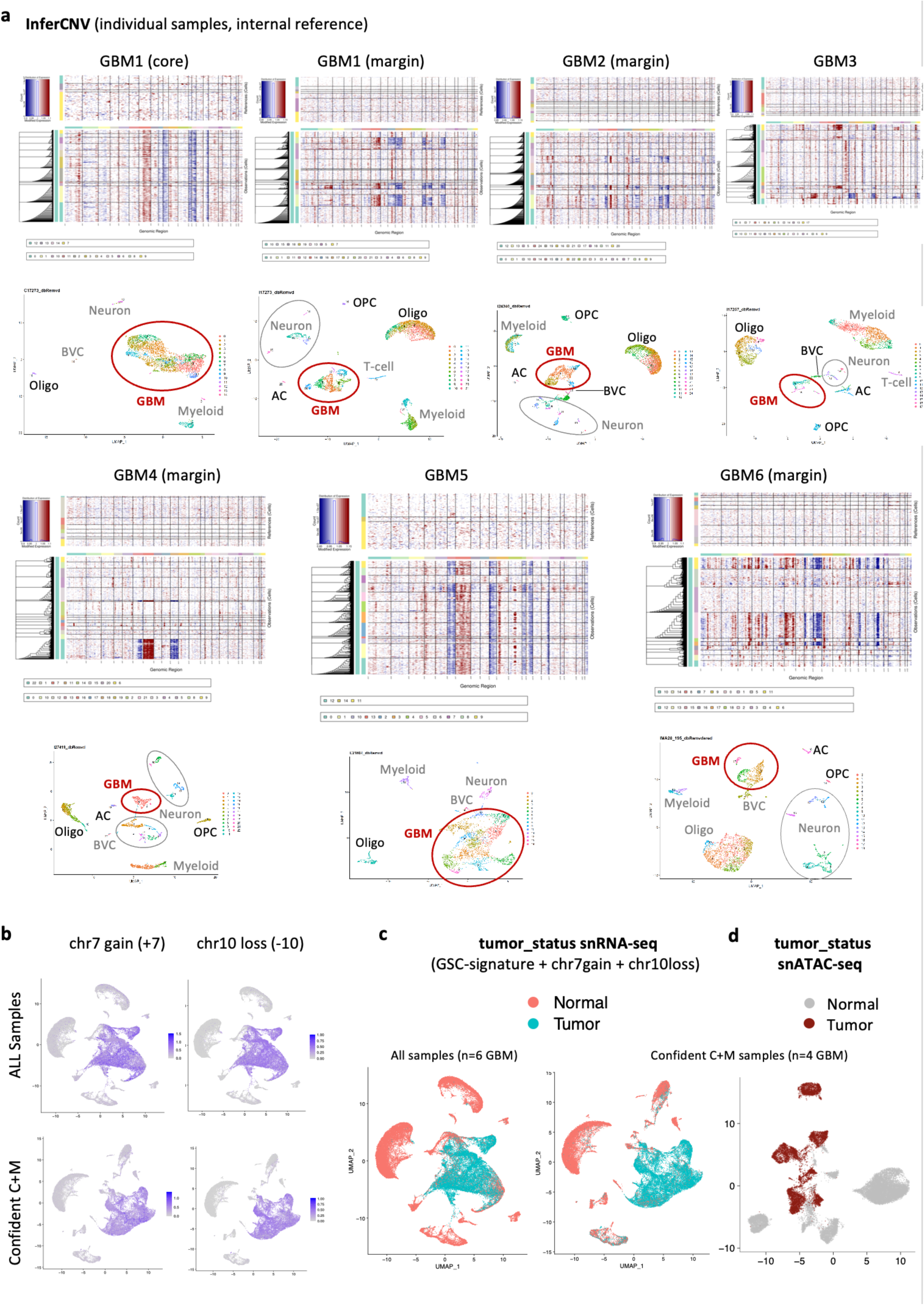
(relates to Fig. 1) **a.** Heatmaps showing inferred copy number variations using InferCNV for each GBM sample using select cell types as internal references (top) with corresponding UMAP cluster-annotated by cell type (bottom). GBM clusters annotated in red; non-neoplastic cell types used as reference annotated in gray; other non-neoplastic cell types annotated in black. **b.** UMAP feature plots showing proportion scaled chr7 gain (+7) or chr 10 loss (-10) inferred from snRNA-seq data in all GBM samples (n=6+6) (top) and for confidently dissected GBM samples (n=4+4) (bottom). **c.** UMAP of integrated snRNA-seq data for all GBM samples (n=6+6) (left) and confidently dissected GBM samples (n=4+4) (right), colored by tumor status. “Tumor_status” is derived from a combined +7, -10, and GSC-signature score. **d.** UMAP of merged snATAC-seq data for GBM samples (n=4+4), colored by RNA-projected tumor status.

**Supplementary Fig. 5:**
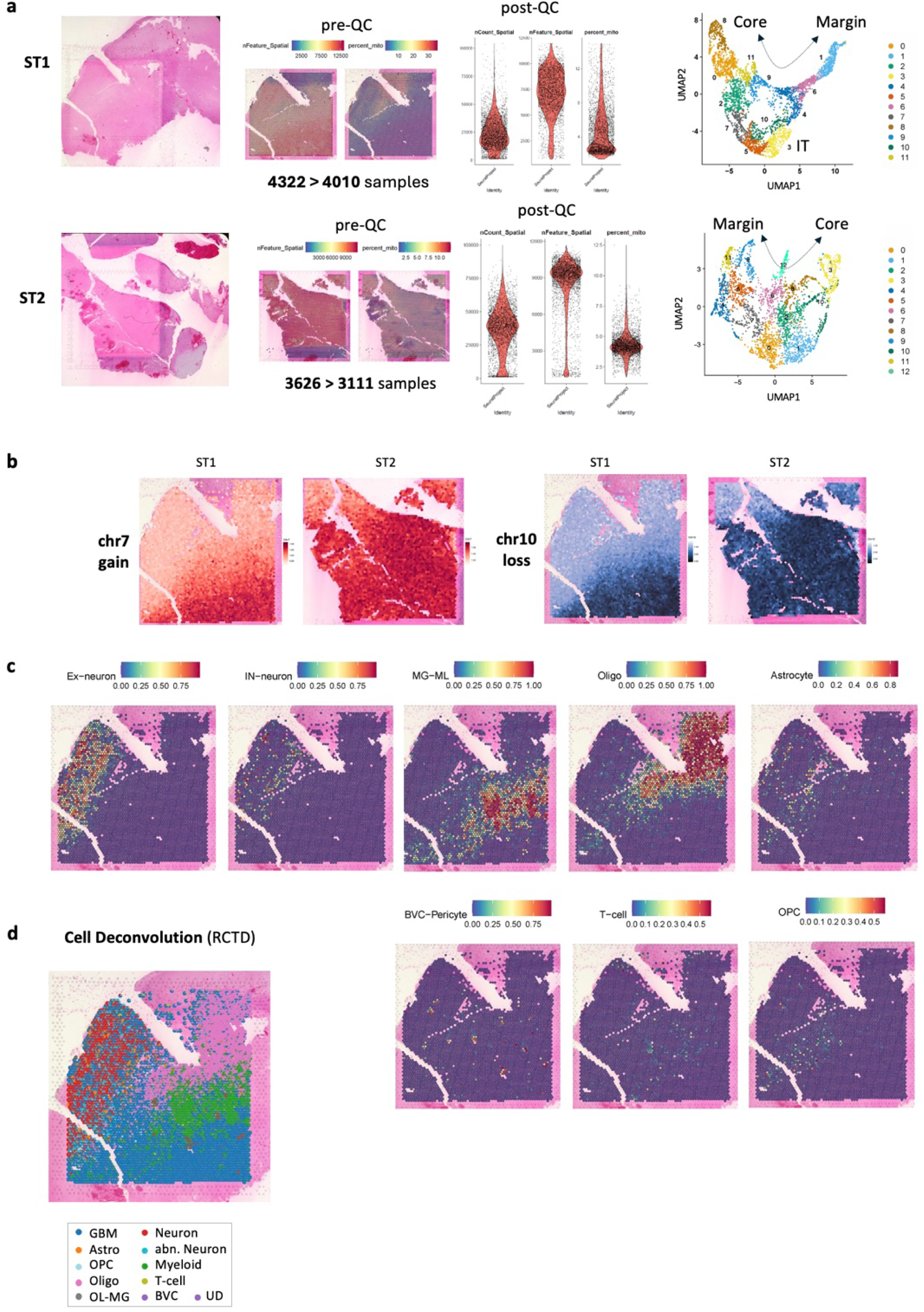
(relates to Fig. 1) **a.** Quality control (QC) of 10x Visium ST1 and ST2 data, showing from left to right: H&E-stained tissue specimens used for spatial analysis; feature plots of features and percent mitochondrial genes pre-QC; violin plots of counts, features, and percent mitochondrial genes post-QC; UMAP embedding colored by cluster number post-QC. **b.** SpatialDimplots showing InferCNV results from SPATA2. Heatmap color denotes chr7gain (red) and chr10loss scores (blue) in ST1 and ST2. **c.** SpatialDimplots of ST1 showing cell type signature projections from snRNA-seq data. **d.** Cell-type deconvolution in ST1 using RCTD showing cell type-enriched ROIs.

**Supplementary Fig. 6.**
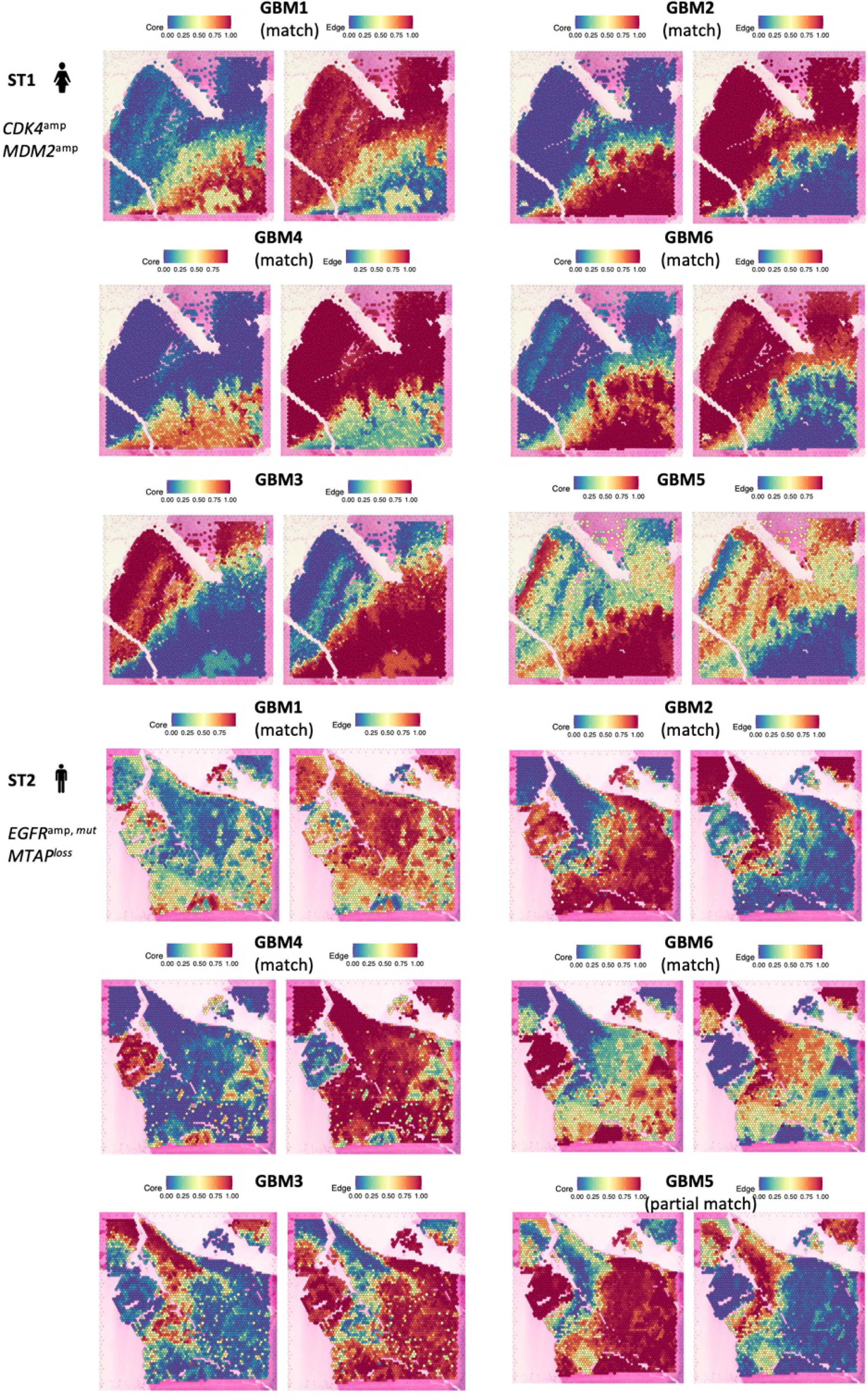
(relates to Fig. 1) SpatialDimplots show projections of individual snRNA-seq GBM patient data from Core and from Margin, onto spatially annotated ST1 and ST2. A match is confirmed for GBM1, GBM2, GBM4, and GBM6.

**Supplementary Fig. 7:**
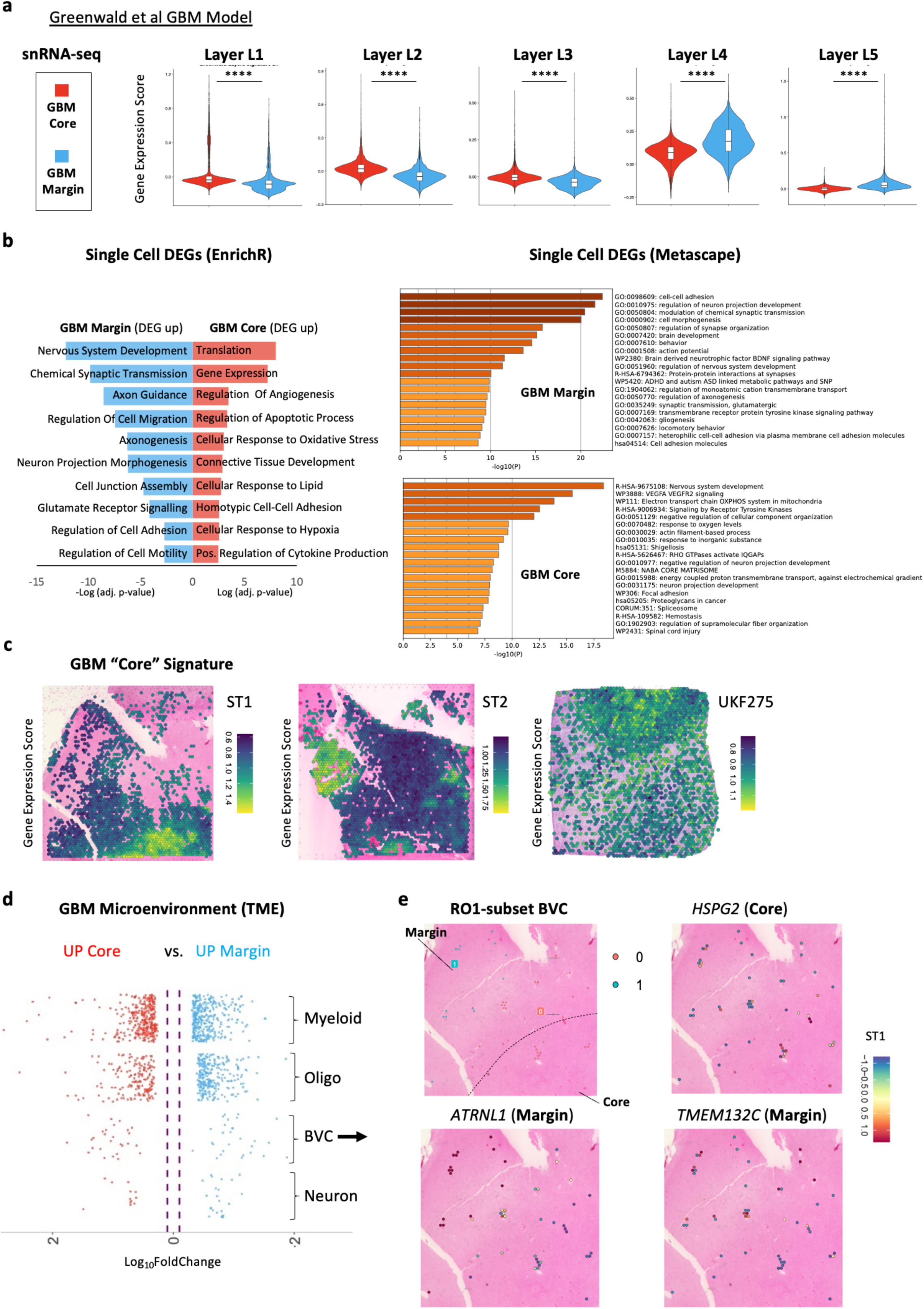
(relates to Fig. 3) **a.** Violin plot of Greenwald et al Layer 1-5 score enrichment in tumor subset GBM-Core and GBM-Margin snRNA-seq populations (n=4+4) (* = p.adj < e^-100^, ** = p.adj < e^-200^, *** = p.adj < e^-300^, **** = p.adj < e^-400^, unpaired Wilcoxon rank test). **b.** Bar plots showing GO term enrichment analyses of GBM-Core and GBM-Margin DEGs from single cell level analysis, using EnrichR (top curated) and Metascape (top 20). **c.** SpatialDimPlot showing enrichment of “GBM core” snRNA-seq signature projected onto ST1, ST2 and UKF275. **d.** Summarized volcano plot showing differentially expressed genes in Core vs. Margin for selected non-malignant (TME) cell types, from confidently dissected snRNA-seq data (n=4+4); (p-adj < 0.05 and log10(FC) > 0.25). Only cell types with >100 cells analyzed. **e.** SpatialDimPlot of subset BVC-enriched ROI showing select markers enriched in core (top) and margin (bottom).

**Supplementary Fig. 8.**
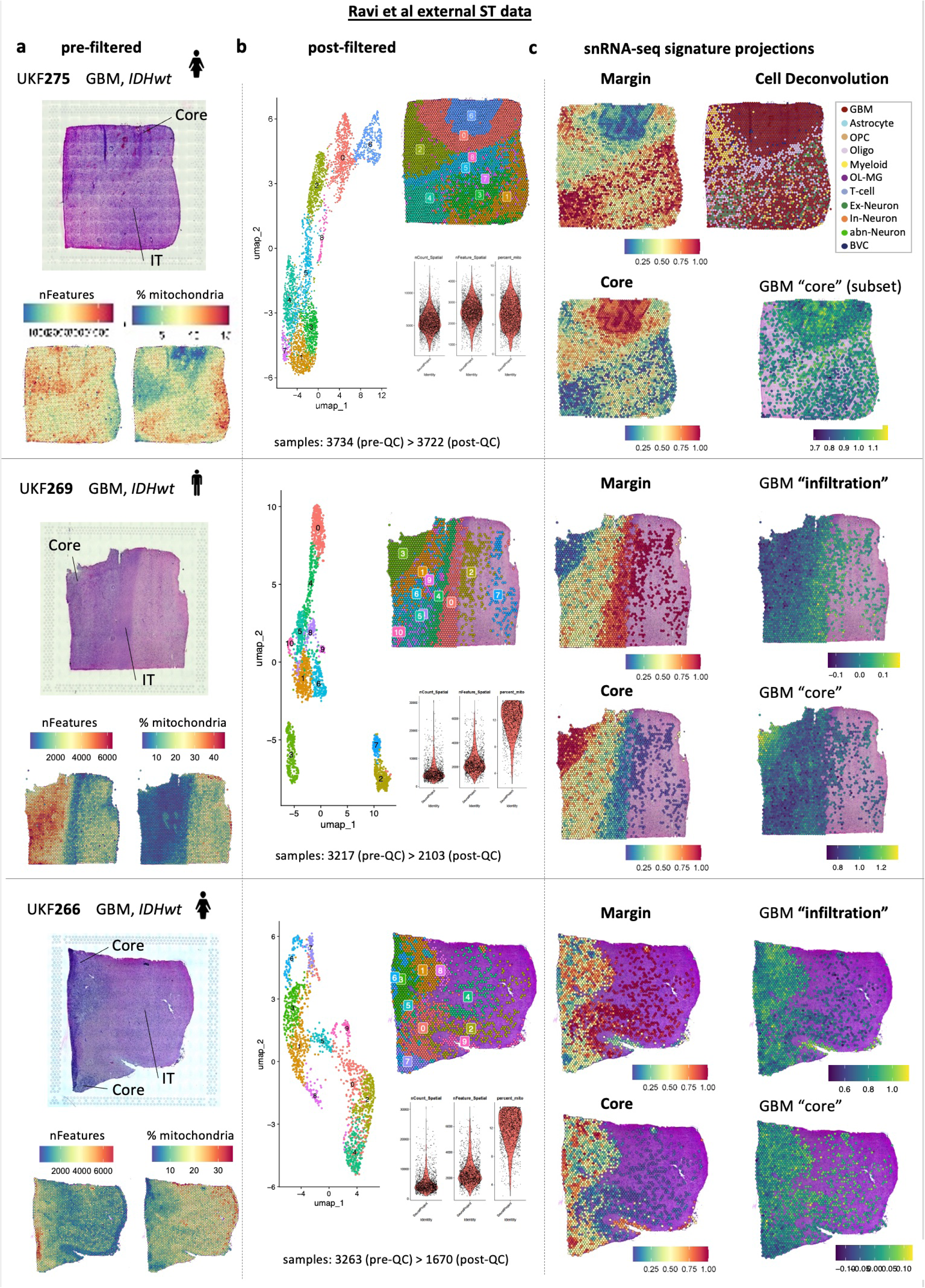
(relates to Fig. 3) 10X Visium analysis of external data (Ravi et al, 2022) for three samples: UKF275 (top), UKF269 (middle) and UKF266 (bottom). **a.** Annotated histological image (top) and SpatialDimPlots (bottom) showing pre-filtered features and percent mitochondrial genes. **b.** Post-filtered data showing UMAP and SpatialDimPlot colored by cluster number, and Violin Plots of QC metrics used. **c.** SpatialDimPlots showing projected scores of snRNA-seq n=4 Margin data (top left), snRNA-seq n=4 Core data (bottom left), “GBM infiltration” signature (top right), “GBM core” signature (bottom right). For the highest quality sample UKF275, cell type predictions from snRNA-seq data are shown instead (top right) and GBM core signature is projected onto GBM-enriched ROI.

**Supplementary Fig. 9:**
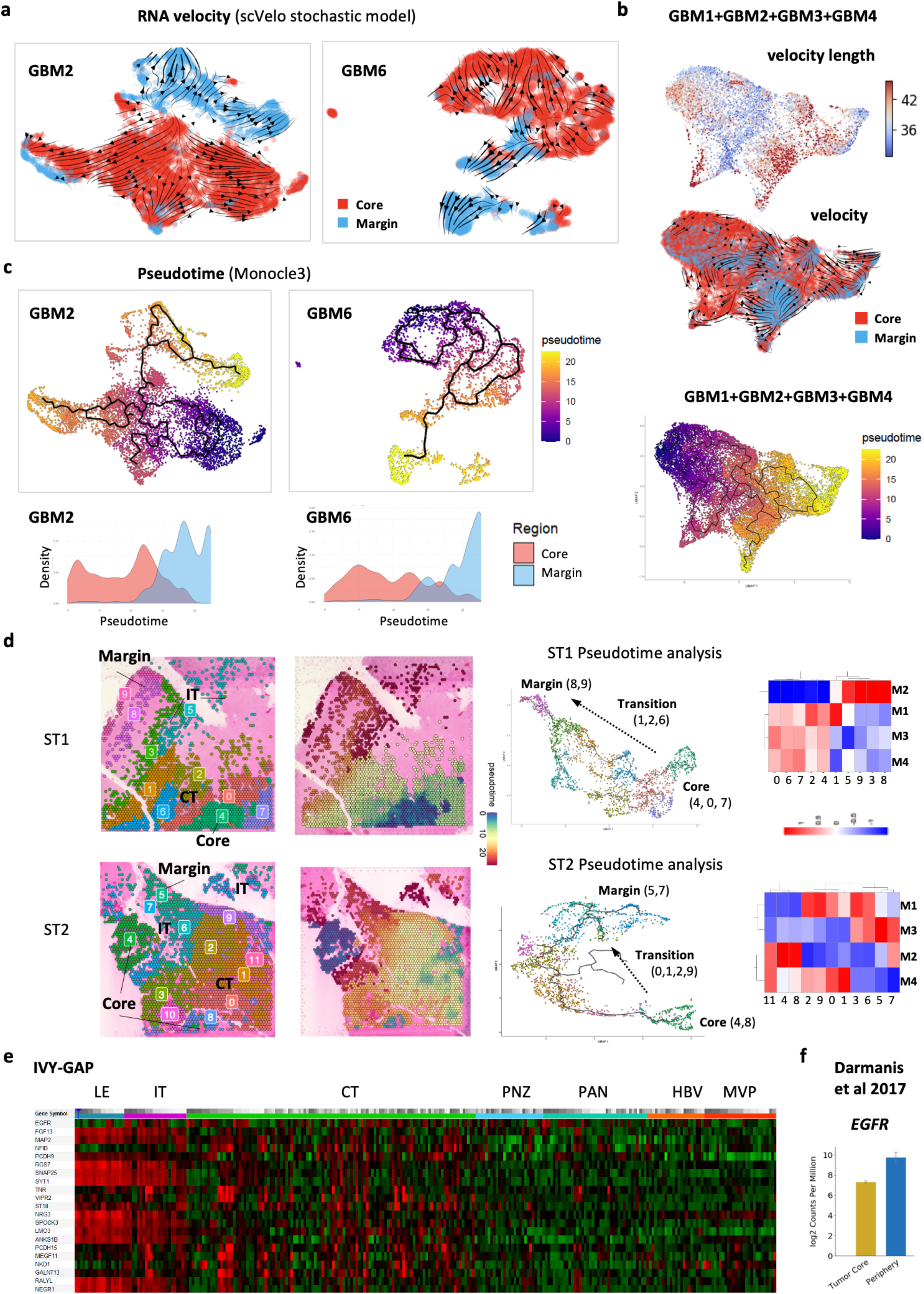
(relates to Fig. 4) **a.** Streamlines of RNA velocity on UMAP, calculated using the stochastic scVelo model, showing predominant core-to-margin directionality for GBM2 (left) and GBM6 (right). **b.** Combined (n=4+4) GBM-subset UMAP embeddings of velocity length (top), RNA velocity (middle), and pseudotime (bottom). **c.** UMAPs (top) and density plots (bottom) of pseudotime, calculated using Monocle3, showing core-to-margin trajectory for GBM2 (left) and GBM6 (right). **d.** Pseudotime analysis using monocle3 in spatial data, ST1(top panels) and ST2 (bottom panels). Shown are SpatialDimPlots colored by cluster number and pseudotime heatmap (left); corresponding core-to-margin UMAPs colored by cluster number (middle); and gene module heatmaps identified across different clusters (right). **e.** Heatmap showing expression of select “GBM infiltration” genes in bulk RNA-seq data from IVY-GAP database across tumor regions. **f.** Bar plot showing expression of *EGFR* in external scRNA-seq dataset of GBM core vs. periphery (Darmanis et al 2017).

**Supplementary Fig. 10:**
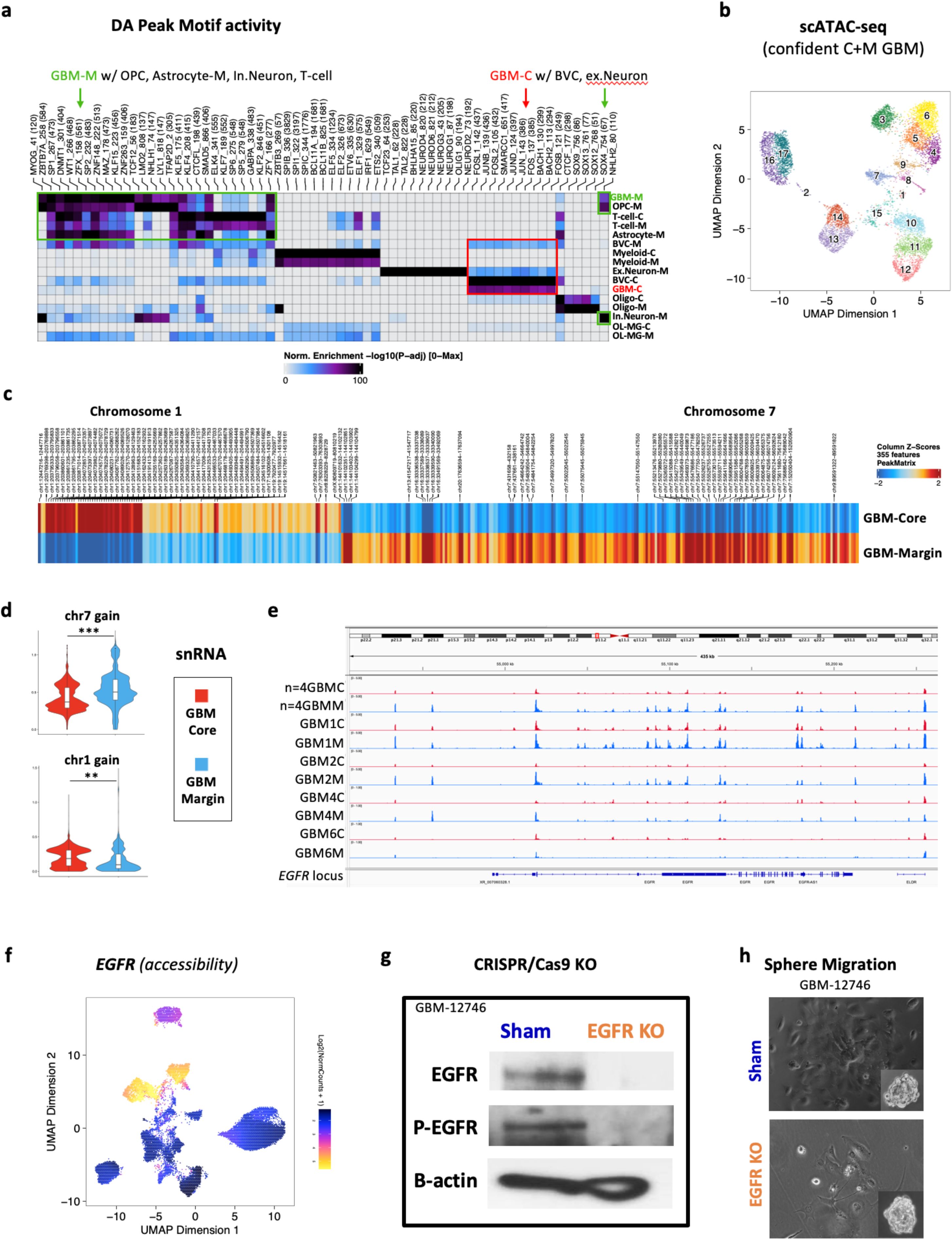
(relates to Fig. 5) **a.** Heatmap showing highest motif activity in differential snATAC-seq peaks, in confidently dissected GBM snATAC-seq data (n=4+4). (FDR < 1, Log2FC >= 0.1, minimal cell number=100). Transcription factors of interest are boxed in green (GBM-Margin shared with OPC, astrocytes, inhibitory neurons, and T-cells) or red (GBM-Core shared with BVC and excitatory Neurons). **b.** UMAP of snATAC-seq subset GBM-Core and GBM-Margin data (n =4+4), colored by cluster number. **c.** Heatmap showing top 50 differential peak locations identified in subset GBM-Core vs. GBM-Margin snATAC-seq data. (FDR < 0.05 & Log2FC >= 0.1 & MeanDiff > 0.2). **d.** Violin plots showing differential enrichment for chr7 gain in GBM-Margin and for chr1 gain in GBM-Core (** = p.adj < e^-200^, *** = p.adj < e^-300^, unpaired Wilcoxon rank test). **e.** IGV plot of snATAC-seq read coverage at EGFR locus, separated by region as well as by sample. **f.** UMAP feature plot showing EGFR accessibility in confidently dissected ATAC-seq data (n=4+4). **g.** Representative immunoblot images showing EGFR deletion and phospho-EGFR (pEGFR) loss after CRISPR/Cas9 knockdown (EGFR KO) in GBM-12746 patient-derived cells, compared to sham. ß-actin used as loading control. **h.** Representative images of spheres at 1h (inset) and 36h in Sham and EGFR KO GBM-12746 cells.

**Supplementary Fig. 11:**
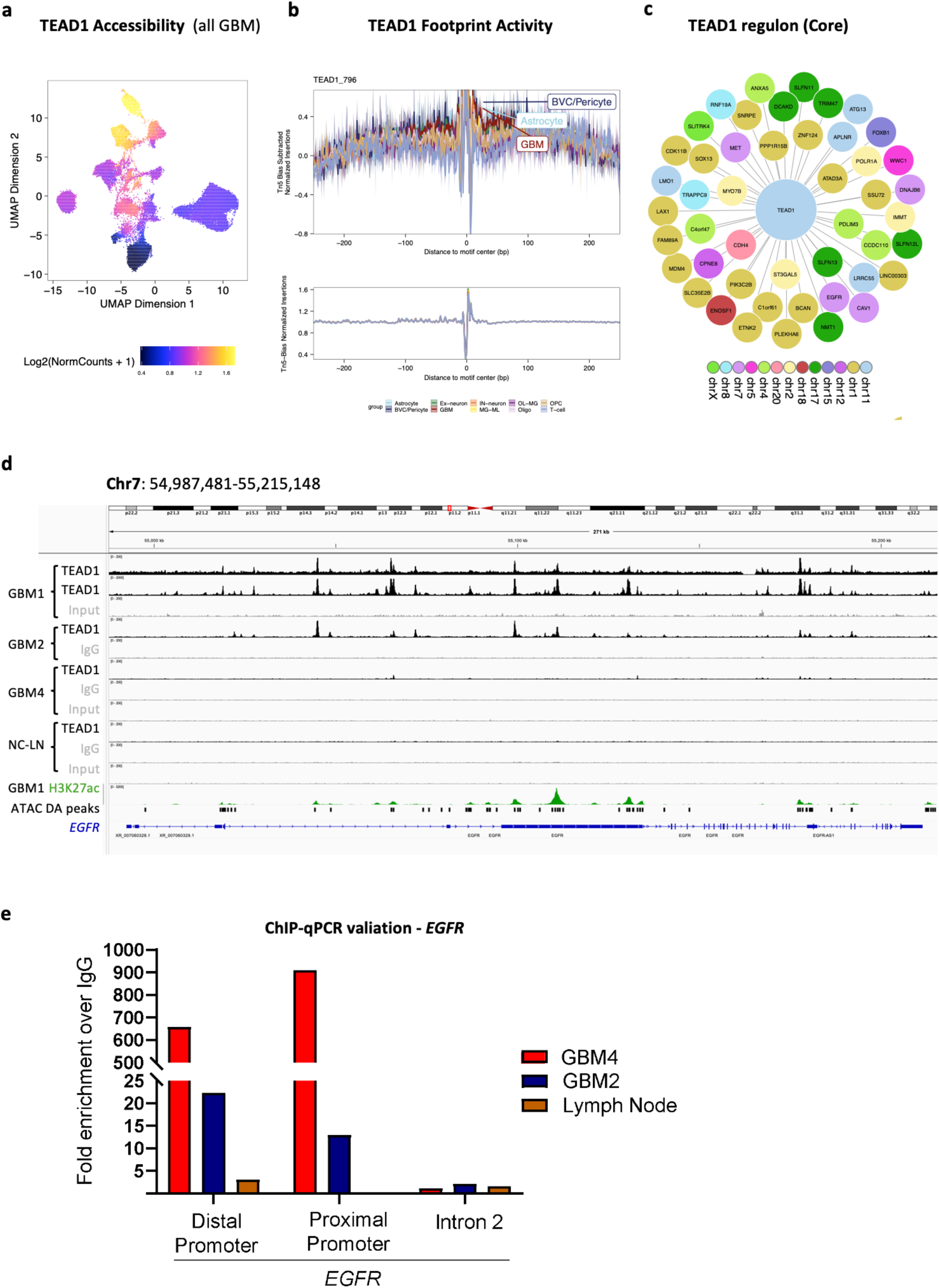
(relates to Fig. 6) **a.** UMAP feature plot showing TEAD1 accessibility in all GBM samples (n=6+6). **b.** TEAD1 footprint profiles across all cell types. GBM, Astrocytes and BVC/Pericyte are labeled. **c.** Regulon analysis of GBM-subset snATAC-seq data showing top TEAD1 target genes in GBM-Core (correlation coefficient > 0.87). **d.** IGV plot of ChIP-seq read coverage at EGFR locus in each sample. Representative pulldowns with TEAD1, H3K27ac, IgG and input are shown. Representative ChIP-qPCR, showing input-normalized Ct enrichment in TEAD1 over IgG pulldown, at *EGFR* distal and proximal promoters in GBM2 and GBM4, compared to EGFR intron2 (internal negative control) and LN (negative tissue control).

## References

1. Tsankova, N.M. and P. Canoll, Advances in genetic and epigenetic analyses of gliomas: a neuropathological perspective. J Neurooncol, 2014. 119(3): p. 481–90.

2. Louis, D.N., et al., The 2021 WHO Classification of Tumors of the Central Nervous System: a summary. Neuro Oncol, 2021. 23(8): p. 1231-1251.

3. Xie, Q., S. Mittal, and M.E. Berens, Targeting adaptive glioblastoma: an overview of proliferation and invasion. Neuro Oncol, 2014. 16(12): p. 1575–84.

4. Schaf, L.R. and I.K. Mellinghof, Glioblastoma and Other Primary Brain Malignancies in Adults: A Review. JAMA, 2023. 329(7): p. 574–587.

5. Gill, B.J., et al., MRI-localized biopsies reveal subtype-specific differences in molecular and cellular composition at the margins of glioblastoma. Proc Natl Acad Sci U S A, 2014. 111(34): p. 12550–5.

6. Hoelzinger, D.B., et al., Gene expression profile of glioblastoma multiforme invasive phenotype points to new therapeutic targets. Neoplasia, 2005. 7(1): p. 7–16.

7. Cooper, L.A., et al., The tumor microenvironment strongly impacts master transcriptional regulators and gene expression class of glioblastoma. Am J Pathol, 2012. 180(5): p. 2108–19.

8. Cuddapah, V.A., et al., A neurocentric perspective on glioma invasion. Nat Rev Neurosci, 2014. 15(7): p. 455–65.

9. Lathia, J.D., et al., Cancer stem cells in glioblastoma. Genes Dev, 2015. 29(12): p. 1203–17.

10. Lan, X., et al., Fate mapping of human glioblastoma reveals an invariant stem cell hierarchy. Nature, 2017. 549(7671): p. 227-232.

11. de Leeuw, C.N. and M.A. Vogelbaum, Supratotal resection in glioma: a systematic review. Neuro Oncol, 2019. 21(2): p. 179–188.

12. Patel, A.P., et al., Single-cell RNA-seq highlights intratumoral heterogeneity in primary glioblastoma. Science, 2014. 344(6190): p. 1396-401.

13. Muller, S., et al., Single-cell profiling of human gliomas reveals macrophage ontogeny as a basis for regional differences in macrophage activation in the tumor microenvironment. Genome Biol, 2017. 18(1): p. 234.

14. Tirosh, I. and M.L. Suva, Dissecting human gliomas by single-cell RNA sequencing. Neuro Oncol, 2018. 20(1): p. 37–43.

15. Yuan, J., et al., Single-cell transcriptome analysis of lineage diversity in high-grade glioma. Genome Med, 2018. 10(1): p. 57.

16. Ne tel, C., et al., An Integrative Model of Cellular States, Plasticity, and Genetics for Glioblastoma. Cell, 2019. 178(4): p. 835–849 e21.

17. Wang, L., et al., The Phenotypes of Proliferating Glioblastoma Cells Reside on a Single Axis of Variation. Cancer Discov, 2019. 9(12): p. 1708–1719.

18. Garo ano, L., et al., Pathway-based classification of glioblastoma uncovers a mitochondrial subtype with therapeutic vulnerabilities. Nat Cancer, 2021. 2(2): p. 141–156.

19. Wang, L., et al., A single-cell atlas of glioblastoma evolution under therapy reveals cell-intrinsic and cell-extrinsic therapeutic targets. Nat Cancer, 2022. 3(12): p. 1534–1552.

20. Liu, M., et al., Spatial transcriptomics reveals segregation of tumor cell states in glioblastoma and marked immunosuppression within the perinecrotic niche. Acta Neuropathol Commun, 2024. 12(1): p. 64.

21. Al-Dalahmah, O., et al., Re-convolving the compositional landscape of primary and recurrent glioblastoma reveals prognostic and targetable tissue states. Nat Commun, 2023. 14(1): p. 2586.

22. Darmanis, S., et al., Single-Cell RNA-Seq Analysis of Infiltrating Neoplastic Cells at the Migrating Front of Human Glioblastoma. Cell Rep, 2017. 21(5): p. 1399–1410.

23. Yu, K., et al., Surveying brain tumor heterogeneity by single-cell RNA-sequencing of multi-sector biopsies. Natl Sci Rev, 2020. 7(8): p. 1306–1318.

24. Venkataramani, V., et al., Glioblastoma hijacks neuronal mechanisms for brain invasion. Cell, 2022. 185(16): p. 2899–2917 e31.

25. Levitin, H.M., et al., De novo gene signature identification from single-cell RNA-seq with hierarchical Poisson factorization. Mol Syst Biol, 2019. 15(2): p. e8557.

26. Aubry, M., et al., From the core to beyond the margin: a genomic picture of glioblastoma intratumor heterogeneity. Oncotarget, 2015. 6(14): p. 12094–109.

27. Patel, K.S., et al., Single-nucleus expression characterization of non-enhancing region of recurrent high-grade glioma. Neurooncol Adv, 2024. 6(1): p. vdae005.

28. Greenwald, A.C., et al., Integrative spatial analysis reveals a multi-layered organization of glioblastoma. Cell, 2024. 187(10): p. 2485–2501 e26.

29. Harwood, D.S.L., et al., Glioblastoma cells increase expression of notch signaling and synaptic genes within infiltrated brain tissue. Nat Commun, 2024. 15(1): p. 7857.

30. Kueckelhaus, J., et al., Inferring histology-associated gene expression gradients in spatial transcriptomic studies. Nat Commun, 2024. 15(1): p. 7280.

31. Ravi, V.M., et al., Spatially resolved multi-omics deciphers bidirectional tumor-host interdependence in glioblastoma. Cancer Cell, 2022. 40(6): p. 639–655 e13.

32. Zheng, Y., et al., Spatial cellular architecture predicts prognosis in glioblastoma. Nat Commun, 2023. 14(1): p. 4122.

33. Manoharan, V.T., et al., Spatiotemporal modeling reveals high-resolution invasion states in glioblastoma. Genome Biol, 2024. 25(1): p. 264.

34. Abdel attah, N., et al., Single-cell analysis of human glioma and immune cells identifies S100A4 as an immunotherapy target. Nat Commun, 2022. 13(1): p. 767.

35. Xie, Y., et al., Key molecular alterations in endothelial cells in human glioblastoma uncovered through single-cell RNA sequencing. JCI Insight, 2021. 6(15).

36. Castellan, M., et al., Single-cell analyses reveal YAP/TAZ as regulators of stemness and cell plasticity in Glioblastoma. Nat Cancer, 2021. 2(2): p. 174–188.

37. Tome-Garcia, J., et al., Analysis of chromatin accessibility uncovers TEAD1 as a regulator of migration in human glioblastoma. Nat Commun, 2018. 9(1): p. 4020.

38. Barrette, A.M., et al., Anti-invasive efficacy and survival benefit of the YAP-TEAD inhibitor verteporfin in preclinical glioblastoma models. Neuro Oncol, 2022. 24(5): p. 694-707.

39. Umphlett, M., et al., IDH-mutant astrocytoma with EGFR amplification-Genomic profiling in four cases and review of literature. Neurooncol Adv, 2022. 4(1): p. vdac067.

40. Richardson, T.E., et al., Genetic and epigenetic instability as an underlying driver of progression and aggressive behavior in IDH-mutant astrocytoma. Acta Neuropathol, 2024. 148(1): p. 5.

41. Ha emeister, C. and R. Satija, Normalization and variance stabilization of single-cell RNA-seq data using regularized negative binomial regression. Genome Biol, 2019. 20(1): p. 296.

42. Hao, Y., et al., Dictionary learning for integrative, multimodal and scalable single-cell analysis. Nat Biotechnol, 2024. 42(2): p. 293–304.

43. Pai, B., et al., High-resolution transcriptomics informs glial pathology in human temporal lobe epilepsy. Acta Neuropathol Commun, 2022. 10(1): p. 149.

44. Jakel, S., et al., Altered human oligodendrocyte heterogeneity in multiple sclerosis. Nature, 2019. 566(7745): p. 543-547.

45. Falcao, A.M., et al., Disease-specific oligodendrocyte lineage cells arise in multiple sclerosis. Nat Med, 2018. 24(12): p. 1837–1844.

46. Ramos, S.I., et al., An atlas of late prenatal human neurodevelopment resolved by single-nucleus transcriptomics. Nat Commun, 2022. 13(1): p. 7671.

47. Granja, J.M., et al., ArchR is a scalable software package for integrative single-cell chromatin accessibility analysis. Nat Genet, 2021. 53(3): p. 403–411.

48. Korsunsky, I., et al., Fast, sensitive and accurate integration of single-cell data with Harmony. Nat Methods, 2019. 16(12): p. 1289–1296.

49. Cable, D.M., et al., Robust decomposition of cell type mixtures in spatial transcriptomics. Nat Biotechnol, 2022. 40(4): p. 517–526.

50. Ribatti, D., R. Tamma, and T. Annese, Epithelial-Mesenchymal Transition in Cancer: A Historical Overview. Transl Oncol, 2020. 13(6): p. 100773.

51. Anastassiou, D., et al., Human cancer cells express Slug-based epithelial-mesenchymal transition gene expression signature obtained in vivo. BMC Cancer, 2011. 11: p. 529.

52. Winkler, F., et al., Cancer neuroscience: State of the field, emerging directions. Cell, 2023. 186(8): p. 1689–1707.

53. Henrik Heiland, D., et al., Tumor-associated reactive astrocytes aid the evolution of immunosuppressive environment in glioblastoma. Nat Commun, 2019. 10(1): p. 2541.

54. Chen, Z., et al., A paracrine circuit of IL-1beta/IL-1R1 between myeloid and tumor cells drives genotype-dependent glioblastoma progression. J Clin Invest, 2023.

55. Bergen, V., et al., Generalizing RNA velocity to transient cell states through dynamical modeling. Nat Biotechnol, 2020. 38(12): p. 1408–1414.

56. La Manno, G., et al., RNA velocity of single cells. Nature, 2018. 560(7719): p. 494-498.

57. Cao, J., et al., A human cell atlas of fetal gene expression. Science, 2020. 370(6518).

58. Puchalski, R.B., et al., An anatomic transcriptional atlas of human glioblastoma. Science, 2018. 360(6389): p. 660-663.

59. Buenrostro, J.D., et al., Single-cell chromatin accessibility reveals principles of regulatory variation. Nature, 2015. 523(7561): p. 486-90.

60. Tchieu, J., et al., NFIA is a gliogenic switch enabling rapid derivation of functional human astrocytes from pluripotent stem cells. Nat Biotechnol, 2019. 37(3): p. 267–275.

61. Brennan, C.W., et al., The somatic genomic landscape of glioblastoma. Cell, 2013. 155(2): p. 462-77.

62. Er ani, P., et al., EGFR promoter exhibits dynamic histone modifications and binding of ASH2L and P300 in human germinal matrix and gliomas. Epigenetics, 2015. 10(6): p. 496–507.

63. Tome-Garcia, J., et al., Prospective Isolation and Comparison of Human Germinal Matrix and Glioblastoma EGFR(+) Populations with Stem Cell Properties. Stem Cell Reports, 2017. 8(5): p. 1421–1429.

64. Reynolds, B.A. and S. Weiss, Clonal and population analyses demonstrate that an EGF-responsive mammalian embryonic CNS precursor is a stem cell. Dev Biol, 1996. 175(1): p. 1–13.

65. Mazzoleni, S., et al., Epidermal growth factor receptor expression identifies functionally and molecularly distinct tumor-initiating cells in human glioblastoma multiforme and is required for gliomagenesis. Cancer Res, 2010. 70(19): p. 7500–13.

66. Guilhamon, P., et al., Single-cell chromatin accessibility profiling of glioblastoma identifies an invasive cancer stem cell population associated with lower survival. Eli e, 2021. 10.

67. Raviram, R., et al., Integrated analysis of single-cell chromatin state and transcriptome identified common vulnerability despite glioblastoma heterogeneity. Proc Natl Acad Sci U S A, 2023. 120(20): p. e2210991120.

68. Wu, S., et al., Circular ecDNA promotes accessible chromatin and high oncogene expression. Nature, 2019. 575(7784): p. 699-703.

69. Zhu, Y., et al., Oncogenic extrachromosomal DNA functions as mobile enhancers to globally amplify chromosomal transcription. Cancer Cell, 2021. 39(5): p. 694–707 e7.

70. Hung, K.L., et al., ecDNA hubs drive cooperative intermolecular oncogene expression. Nature, 2021. 600(7890): p. 731-736.

71. Snuderl, M., et al., Mosaic amplification of multiple receptor tyrosine kinase genes in glioblastoma. Cancer Cell, 2011. 20(6): p. 810–7.

72. Okada, Y., et al., Selection pressures of TP53 mutation and microenvironmental location influence epidermal growth factor receptor gene amplification in human glioblastomas. Cancer Res, 2003. 63(2): p. 413–6.

73. Talasila, K.M., et al., EGFR wild-type amplification and activation promote invasion and development of glioblastoma independent of angiogenesis. Acta Neuropathol, 2013. 125(5): p. 683–98.

74. Parker, J.J., et al., Gefitinib selectively inhibits tumor cell migration in EGFR-amplified human glioblastoma. Neuro Oncol, 2013. 15(8): p. 1048–57.

75. Guillamo, J.S., et al., Molecular mechanisms underlying effects of epidermal growth factor receptor inhibition on invasion, proliferation, and angiogenesis in experimental glioma. Clin Cancer Res, 2009. 15(11): p. 3697–704.

76. Lund-Johansen, M., et al., Effect of epidermal growth factor on glioma cell growth, migration, and invasion in vitro. Cancer Res, 1990. 50(18): p. 6039–44.

77. Reardon, D.A., P.Y. Wen, and I.K. Mellinghof, Targeted molecular therapies against epidermal growth factor receptor: past experiences and challenges. Neuro Oncol, 2014. 16 **Suppl 8**(Suppl 8): p. viii7-13.

78. Masliantsev, K., et al., Yes-Associated Protein Nuclear Translocation Is Regulated by Epidermal Growth Factor Receptor Activation Through Phosphatase and Tensin Homolog/AKT Axis in Glioblastomas. Lab Invest, 2023. 103(5): p. 100053.

79. Okamoto, K., et al., AXL activates YAP through the EGFR-LATS1/2 axis and confers resistance to EGFR-targeted drugs in head and neck squamous cell carcinoma. Oncogene, 2023. 42(39): p. 2869–2877.

80. Gao, M., et al., EGFR Activates a TAZ-Driven Oncogenic Program in Glioblastoma. Cancer Res, 2021. 81(13): p. 3580–3592.

81. Vigneswaran, K., et al., YAP/TAZ Transcriptional Coactivators Create Therapeutic Vulnerability to Verteporfin in EGFR-mutant Glioblastoma. Clin Cancer Res, 2021. 27(5): p. 1553–1569.

82. Read, R.D., Repurposing the drug verteporfin as anti-neoplastic therapy for glioblastoma. Neuro Oncol, 2022. 24(5): p. 708–710.

83. Whitfield, M.L., et al., Identification of genes periodically expressed in the human cell cycle and their expression in tumors. Mol Biol Cell, 2002. 13(6): p. 1977–2000.

84. Zhang, Y., et al., Model-based analysis of ChIP-Seq (MACS). Genome Biol, 2008. 9(9): p. R137.

85. Weirauch, M.T., et al., Determination and inference of eukaryotic transcription factor sequence specificity. Cell, 2014. 158(6): p. 1431–1443.

86. Schep, A.N., et al., chromVAR: inferring transcription-factor-associated accessibility from single-cell epigenomic data. Nat Methods, 2017. 14(10): p. 975–978.

87. Kuleshov, M.V., et al., Enrichr: a comprehensive gene set enrichment analysis web server 2016 update. Nucleic Acids Res, 2016. 44(W1): p. W90–7.

88. Zanconato, F., et al., Genome-wide association between YAP/TAZ/TEAD and AP-1 at enhancers drives oncogenic growth. Nat Cell Biol, 2015. 17(9): p. 1218–27.

